# Quantitative genetic analysis of interactions in the pepper-*Phytophthora capsici* pathosystem

**DOI:** 10.1101/2021.12.05.471294

**Authors:** Gregory Vogel, Garrett Giles, Kelly R. Robbins, Michael A. Gore, Christine D. Smart

## Abstract

The development of pepper cultivars with durable resistance to the oomycete *Phytophthora capsici* has been challenging due to differential interactions between the species that allow certain pathogen isolates to cause disease on otherwise resistant host genotypes. Currently, little is known about the pathogen genes that are involved in these interactions. To investigate the genetic basis of *P. capsici* virulence on individual pepper genotypes, we inoculated sixteen pepper accessions – representing commercial varieties, sources of resistance, and host differentials – with 117 isolates of *P. capsici*, for a total of 1,864 host-pathogen combinations. Analysis of disease outcomes revealed a significant effect of inter-species genotype-by-genotype interactions, although these interactions were quantitative rather than qualitative in scale. Isolates were classified into five pathogen subpopulations, as determined by their genotypes at over 60,000 single-nucleotide polymorphisms (SNPs). While absolute virulence levels on certain pepper accessions significantly differed between subpopulations, a multivariate phenotype reflecting relative virulence levels on certain pepper genotypes compared to others showed the strongest association with pathogen subpopulation. A genome-wide association study (GWAS) identified four pathogen loci significantly associated with virulence, two of which colocalized with putative RXLR effector genes and another with a polygalacturonase gene cluster. All four loci appeared to represent broad-spectrum virulence genes, as significant SNPs demonstrated consistent effects regardless of the host genotype tested. Host genotype-specific virulence variants in *P. capsici* may be difficult to map via GWAS, perhaps controlled by many genes of small effect or by multiple alleles that have arisen independently at the same loci.

## INTRODUCTION

Populations of plant pathogens show both quantitative and qualitative variation for the level of disease severity they cause on host plants. Qualitative differences in a pathogen’s ability to cause disease, or its pathogenicity, are often due to polymorphisms in effector genes that enable the pathogen to evade recognition by a cognate resistance (R) gene in the host. In the classic gene-for-gene model, alleles at each of a single R and effector gene interact to determine the presence or absence of disease (Flor 1955). On the other hand, virulence, or aggressiveness, is commonly defined as the quantitative extent of disease caused by a pathogen (Sacristán and García-Arenal 2008). Variation in virulence – and its counterpart in plants, quantitative disease resistance – is thought to be under polygenic control, governed by many genes of small effect (Nelson et al. 2018; Pariaud et al. 2009). Although it was once proposed that virulence and quantitative disease resistance were broad-spectrum in their effects (Van der Plank 1968), numerous examples of quantitative interactions between host and pathogen have since been described (Andrivon et al. 2007; Parlevliet 1976; Salvaudon et al. 2007). Recent genetic mapping and association studies in both plant and pathogen populations have identified pathogen virulence loci whose effects are conditional on host genotype (Hartmann et al. 2017; Meile et al. 2018; Soltis et al. 2019; Stewart et al. 2018), and conversely, quantitative disease resistance loci in plants whose effects are conditional on pathogen isolate (Corwin et al. 2016; Qi et al. 1999; Wang et al. 2018; Zenbayashi-Sawata et al. 2005). Nevertheless, the genetic basis of quantitative host-pathogen interactions remains poorly understood in most pathosystems.

*Phytophthora capsici* is a highly destructive, oomycete pathogen that causes root, crown, and fruit rots on pepper (*Capsicum annuum*) and several other economically important host species in the Solanaceae, Cucurbitaceae, and Fabaceae families (Granke et al. 2012). As a hemibiotroph, *P. capsici* infects plants without initially causing noticeable disease symptoms, before rapidly transitioning to a necrotrophic phase of disease associated with substantial tissue death (Lamour et al. 2012b). As in other oomycete pathogens, the early, biotrophic stage of disease caused by *P. capsici* is believed to be facilitated by the secretion of hundreds of effector proteins that suppress plant defense responses or play additional roles in virulence (Jupe et al. 2013; Schornack et al. 2009). Effector genes in many species have been shown to condition avirulence when their gene product is recognized by an R protein in the host (Schornack et al. 2009). Two families of effectors have been characterized in oomycetes: RXLRs (Morgan and Kamoun 2007) and ‘Crinklers’ (CRNs) (Torto et al. 2003), which are represented, respectively, by 516 and 84 predicted genes in the *P. capsici* genome (Jupe et al. 2013; Stam et al. 2013b). Virulence activities have been functionally verified for a number of these genes in *P. capsici* (Chen et al. 2019; Fan et al. 2018; Li et al. 2019a, 2019b, 2020; Mafurah et al. 2015; Stam et al. 2013a).

The genetics of resistance to *P. capsici* in pepper have been characterized extensively. Several sources of resistance have been described, including serrano pepper landrace Criollos de Morelos 334 (CM334; Guerrero-Moreno and Laborde 1980) and hot pepper landraces Perennial (Lefebvre and Palloix 1996) and PI 201234 (Kimble and Grogan 1960). These accessions, and perhaps others, have been used commercially to breed resistant cultivars representing several different market classes of pepper. Mapping experiments using crosses with these sources of resistance have identified a major quantitative trait locus (QTL) on chromosome 5 of pepper that is shared by multiple resistant accessions and appears to demonstrate a consistent effect against a broad range of isolates (Mallard et al. 2013; Ogundiwin et al. 2005; Rehrig et al. 2014; Siddique et al. 2019; Thabuis et al. 2003; Truong et al. 2012). In addition, numerous other loci of smaller effect have been mapped that demonstrate evidence of having isolate-specific effects (Ogundiwin et al. 2005; Rehrig et al. 2014; Siddique et al. 2019; Truong et al. 2012).

As expected due to the existence of isolate-specific resistance in the host, *P. capsici* isolates vary in their ability to cause disease on different host genotypes. Certain pathogen isolates, for example, are able to overcome the resistance found in many commercial varieties (Foster and Hausbeck 2010; Parada-Rojas and Quesada-Ocampo 2019). *Phytophthora capsici* isolates have been classified into distinct physiological races using several sets of differential host lines, including the New Mexico Recombinant Inbred Line (NMRIL) population, a set of RILs derived from a cross between CM334 and jalapeño variety Early Jalapeño (Hu et al. 2013; Glosier et al. 2008; Oelke et al. 2003; Sy et al. 2008). These experiments have resulted in race designations that are difficult to translate between populations because of the different lines used in each experiment and the large number of races identified by many researchers, who in several cases have demonstrated that every isolate assayed in an experiment belongs to a separate race (Barchenger et al. 2018b; Monroy-Barbosa and Bosland 2011; Reyes-Tena et al. 2019). Given the complexity of the interactions between pepper and *P. capsici*, and our limited understanding of the genes involved in these interactions, the standardization of race-typing protocols is a major challenge. Further knowledge of the genes underlying virulence variation in *P. capsici*, which remains largely unexplored compared to our understanding of the genetics of resistance in pepper, is needed.

Genetic studies using controlled crosses are possible in *P. capsici*, as recombinant oospores are produced readily in culture when isolates of opposite mating type are paired (Hurtado-Gonzalez and Lamour 2009). Such approaches have been used successfully to determine the inheritance of traits such as fungicide sensitivity (Lamour and Hausbeck 2000) and pathogenicity on different host species (Polach and Webster 1972), but they can be challenging due to technical factors such as labor-intensive protocols for obtaining single-oospore cultures (Hurtado-Gonzalez and Lamour 2009) or biological factors such as frequent mitotic loss of heterozygosity in parental and progeny strains (Carlson et al. 2017; Lamour et al. 2012a). The increased affordability of genome-wide markers obtained by next-generation sequencing have made genome-wide association studies (GWAS) an attractive alternative for exploring the genetic bases of traits in fungal and oomycete pathogens (Bartoli and Roux 2017; Plissonneau et al. 2017). Genome-wide association studies rely on historical recombination events to generate short segments of linkage disequilibrium (LD) blocks across the genome. As a result, in pathogens that undergo frequent sexual reproduction, traits can be mapped with high resolution using natural collections of isolates collected from the field. GWAS have been used to identify both qualitative and quantitative virulence loci in various fungal pathogens, including *Parastagonospora nodorum* (Gao et al. 2016)*, Zymoseptoria tritici* (Hartmann et al. 2017; Zhong et al. 2017), and *Botrytis cinerea* (Soltis et al. 2019).

Recently, we used genotyping-by-sequencing (GBS) to characterize genetic variation in 245 isolates of *P. capsici*, collected largely in New York (NY) state, at over 60,000 single-nucleotide polymorphism (SNP) loci (Vogel et al. 2021). Using a subset of 129 genetically distinct (i.e. non-clonal) isolates, we found evidence for limited gene flow between pathogen subpopulations located on different farms, and discovered loci via GWAS associated with mating type and fungicide sensitivity (Vogel et al. 2021). In the current study, we phenotyped 117 isolates from this genotyped panel for their virulence on each of 16 pepper accessions, with the goal of shedding light on the genetic architecture of virulence in *P. capsici* on pepper. Our specific objectives were to: i) quantify and characterize the nature of genotype-by-genotype interactions between pepper and *P. capsici*; ii) determine the degree of differentiation in virulence phenotypes between pathogen subpopulations collected on different farms; and iii) conduct a GWAS to identify variants in *P. capsici* associated with virulence on one or multiple pepper accessions.

## RESULTS

### Host-isolate interactions in the pepper-*P. capsici* pathosystem

Seedlings of 16 pepper accessions were inoculated in a greenhouse experiment with each of 117 genetically distinct isolates of *P. capsici*, for a total of 2,784 measurements of mortality incidence-based Area Under the Disease Progress Curve (AUDPC), representing 1,864 distinct host × pathogen combinations. Pepper accessions chosen for inclusion in this study represented either sources of disease resistance (CM334 and Perennial), lines used in previous publications as differential hosts (NMRIL-A, NMRIL-G, NMRIL-H, NMRIL-I, NMRIL-N, NMRIL-Z, and Early Jalapeño), or commercial bell pepper hybrids (Archimedes, Aristotle, Intruder, Paladin, Red Knight, Revolution, and Vanguard) featuring a range of overall resistance levels and developed for the eastern U.S., where the majority of the pathogen isolates featured in this study were collected. Inoculations were performed in incomplete blocks with replicated checks, in order to control for batch effects (see Materials and Methods). Two complete replicates of the experiment were conducted, although a subset of pepper accessions that demonstrated complete resistance to all isolates in the first replicate were replaced with different accessions in the second replicate. As a result, each host × pathogen combination was replicated twice for eight of the pepper accessions and once for the other eight.

Of the 117 isolates, 12 failed to cause disease on a single pepper accession in either replicate of the experiment. These non-pathogenic isolates were disproportionately represented by a single field population (Ontario #1) from 2013 that was maintained in long-term storage for longer than the majority of the isolates in this experiment, and the 12 isolates were subsequently excluded from analysis. Of the 16 pepper accessions, eight were completely resistant to over 80% of the remaining isolates (Figure S1). These accessions – comprised of resistant landrace CM334, hybrid bell pepper Intruder, and the six NMRIL differential lines – were excluded from analysis as well, leaving a dataset consisting of 105 isolates and eight pepper accessions: hot pepper landrace Perennial, open-pollinated variety Early Jalapeño, and six hybrid bell pepper cultivars (Archimedes, Aristotle, Paladin, Red Knight, Revolution, and Vanguard).

Mixed linear models were fitted to estimate the effects of pepper accession, pathogen isolate, and accession-isolate interaction on AUDPC. Analysis of variance (ANOVA) in a balanced dataset that excluded three pepper accessions (Archimedes, Revolution, and Vanguard, each included in a single replicate) revealed significant effects for pepper accession (*P* = 2.5 × 10^-32^), *P. capsici* isolate (*P* = 1.3 × 10^-225^), and their interaction (*P* = 4.5 × 10^-49^) (Table 1). Of the experimental design terms included in the model, block (*P* = 4.7 × 10^-7^) and tray (i.e. whole plot error) (*P* = 1.5 × 10^-12^) had significant effects on AUDPC, whereas replicate (*P* = 0.44) did not (Table 1; Table S1). Block and tray collectively accounted for 36.03% of the variance explained by random terms in the model. Broad-sense heritability estimates for virulence on each pepper accession, as estimated in nested models fitted separately for each accession, were moderate to high, ranging from 0.72 for Early Jalapeño to 0.93 for Red Knight (Table 2), indicating that isolate virulence estimates were largely consistent between experimental replicates.

**Table 1:** Test statistics and P-values associated with fixed effects in model testing the effects of isolate, pepper accession, and their interaction on AUDPC.

|  | Df | Denominator DF | F statistic | *P*-value |
| --- | --- | --- | --- | --- |
| Isolate | 107 | 119.5 | 11.07 | $2.50 \times 10^{-32}$ |
| Pepper | 4 | 516 | 851.3 | $1.30 \times 10^{-225}$ |
| Isolate × Pepper | 428 | 516 | 3.98 | $4.50 \times 10^{-49}$ |
| Rep | 1 | 7 | 0.66 | 0.44 |

**Table 2:** Medians, broad-sense heritabilities, and standard errors of heritability for AUDPC on

| Pepper | Median AUDPC | H<sup>2</sup><sup>a</sup> | SE<sup>a</sup> |
| --- | --- | --- | --- |
| Archimedes | 3.27 | NA | NA |
| Aristotle | 23.18 | 0.86 | 0.03 |
| Early Jalapeño | 2.55 | 0.72 | 0.05 |
| Paladin | 2.02 | 0.8 | 0.04 |
| Perennial | 10.74 | 0.79 | 0.04 |
| Red Knight | 52.8 | 0.93 | 0.01 |
| Revolution | 9.49 | NA | NA |
| Vanguard | 8.34 | NA | NA |

<sup>a</sup> Heritabilities and standard errors are not shown for pepper accessions only included in one replicate.

Model-adjusted AUDPC means (i.e., least squares means) were estimated for each host-pathogen combination using a linear model fitted with the whole dataset, including the three pepper accessions included in only one replicate. Virulence, as reflected by AUDPC LS mean, was non-normally distributed on each of the eight pepper accessions (Figure 1). On Red Knight, which was on average the most susceptible of the eight pepper accessions with a median AUDPC of 52.8 (Table 2), virulence had a bimodal distribution, with the majority of isolates able to cause high levels of disease but a smaller, yet substantial, subset of isolates causing low plant mortality. On the remaining pepper accessions, virulence distributions had a mode closer to zero, although they featured heavy right tails, especially in the case of Aristotle, for which virulence was almost uniformly distributed. Early Jalapeño, Paladin, and Archimedes were the most resistant pepper accessions overall (median AUDPCs of 2.55, 2.02, and 3.27, respectively), and while they still featured skewed AUDPC distributions, they had less dense right tails, with only a small number of outlier isolates able to cause high levels of disease. Hierarchical clustering of pepper accessions in terms of similarity in their resistance levels to the 105 isolates showed that accessions did not group based on their market class or improvement status, as hot pepper landrace Perennial clustered with bell pepper hybrids Aristotle, Revolution and Vanguard, and similarly, Early Jalapeño clustered with bell pepper hybrids Paladin and Archimedes (Figure 1).

**Figure 1.**
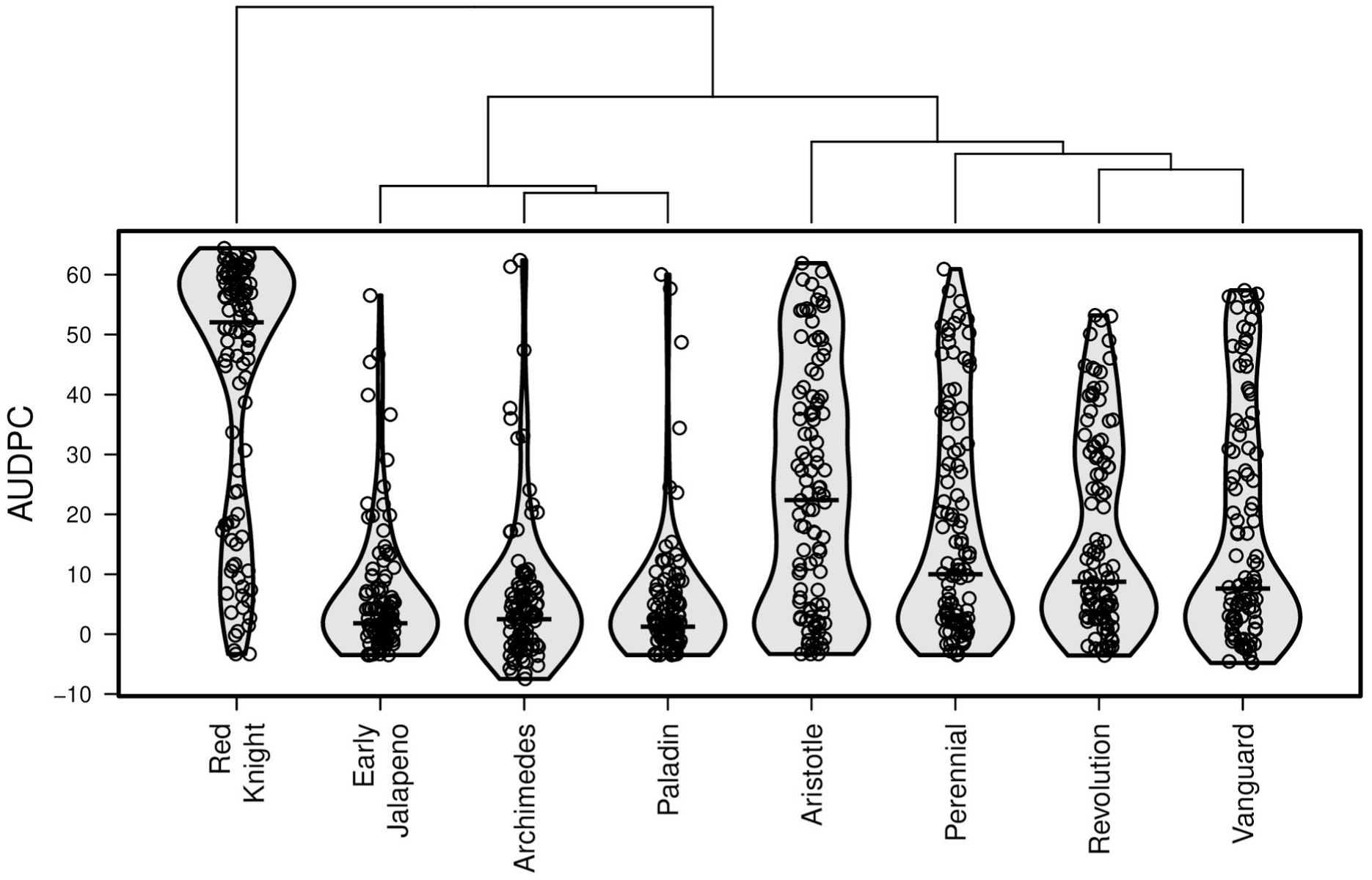
Distributions of AUDPC LS means among 105 isolates on each of 8 pepper accessions. Dendrogram created by hierarchical clustering of peppers based on the Euclidian distance between their AUDPC values for each isolate.

Visualization of the virulence profiles of the 105 isolates on each pepper revealed a diversity of phenotypic patterns (Figure 2). In general, the disease caused by individual isolates tended to be more severe on pepper accessions that were more susceptible on average. However, many exceptions were observed where peppers differed in rank in terms of their susceptibility to individual isolates compared to their overall susceptibility across isolates.

**Figure 2:**
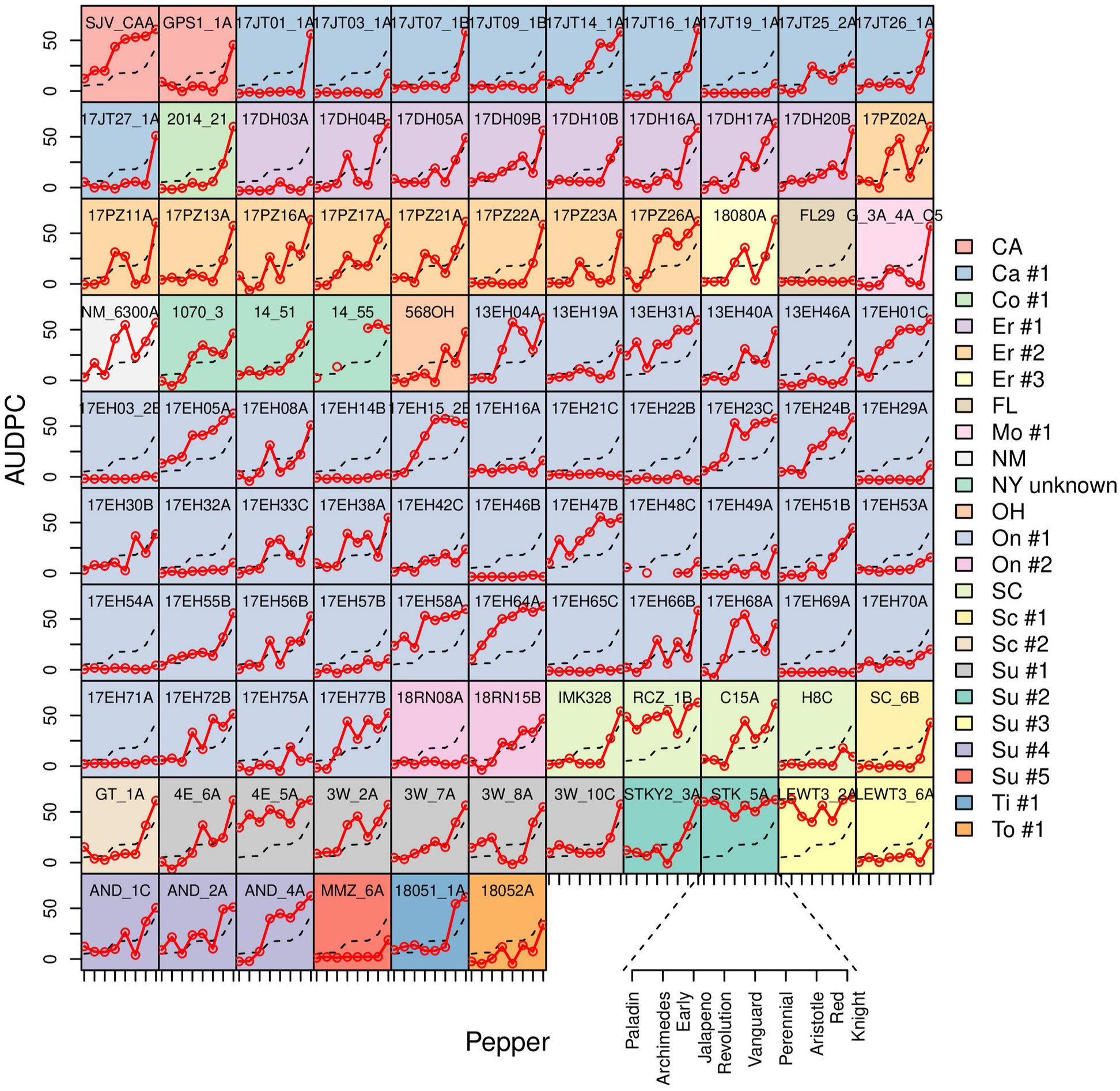
Isolate virulence profiles. Each subplot shows the AUDPC LS mean for a particular isolate on each of 8 pepper accessions in red. The black dashed line signifies the mean AUDPC for pepper, averaged across isolates. Pepper accessions are in order of increasing average susceptibility. Each background color refers to a distinct field site, or state if collected outside NY. CA=California; Ca=Cayuga County, NY; Co=Columbia County, NY; Er=Erie County, NY; FL=Florida; Mo=Monroe County, NY; NM=New Mexico; OH=Ohio; On=Ontario County, NY; SC=South Carolina; Sc=Schenectady County, NY; Su=Suffolk County, NY; Ti=Tioga County, NY; To=Tompkins County, NY.

We explored the relationship between individual isolate virulence profiles and average pepper susceptibility levels by regressing the AUDPC associated with each host-pathogen combination on the average AUDPC for pepper accession, akin to Finlay-Wilkinson analysis, a technique used for characterizing the yield stability of crop varieties in different environments. Isolates fell into one of several categories based on their regression slopes and intercepts (Figure 3). The majority of regression lines had a low intercept and a slope between 1 and 1.5, corresponding to isolates with average or higher than average virulence on more susceptible pepper accessions and average or lower than average virulence on more resistant pepper accessions. Another subset of regression lines had both intercept and slope close to 0, reflecting isolates with consistently low virulence across all of the pepper accessions in the experiment. Finally, the regression lines for four isolates (LEWT3_2A, STK_5A, RCZ_1B, and 4E_5A) had a low slope but a high intercept. These were the only isolates observed to be highly virulent on all eight pepper accessions.

**Figure 3:**
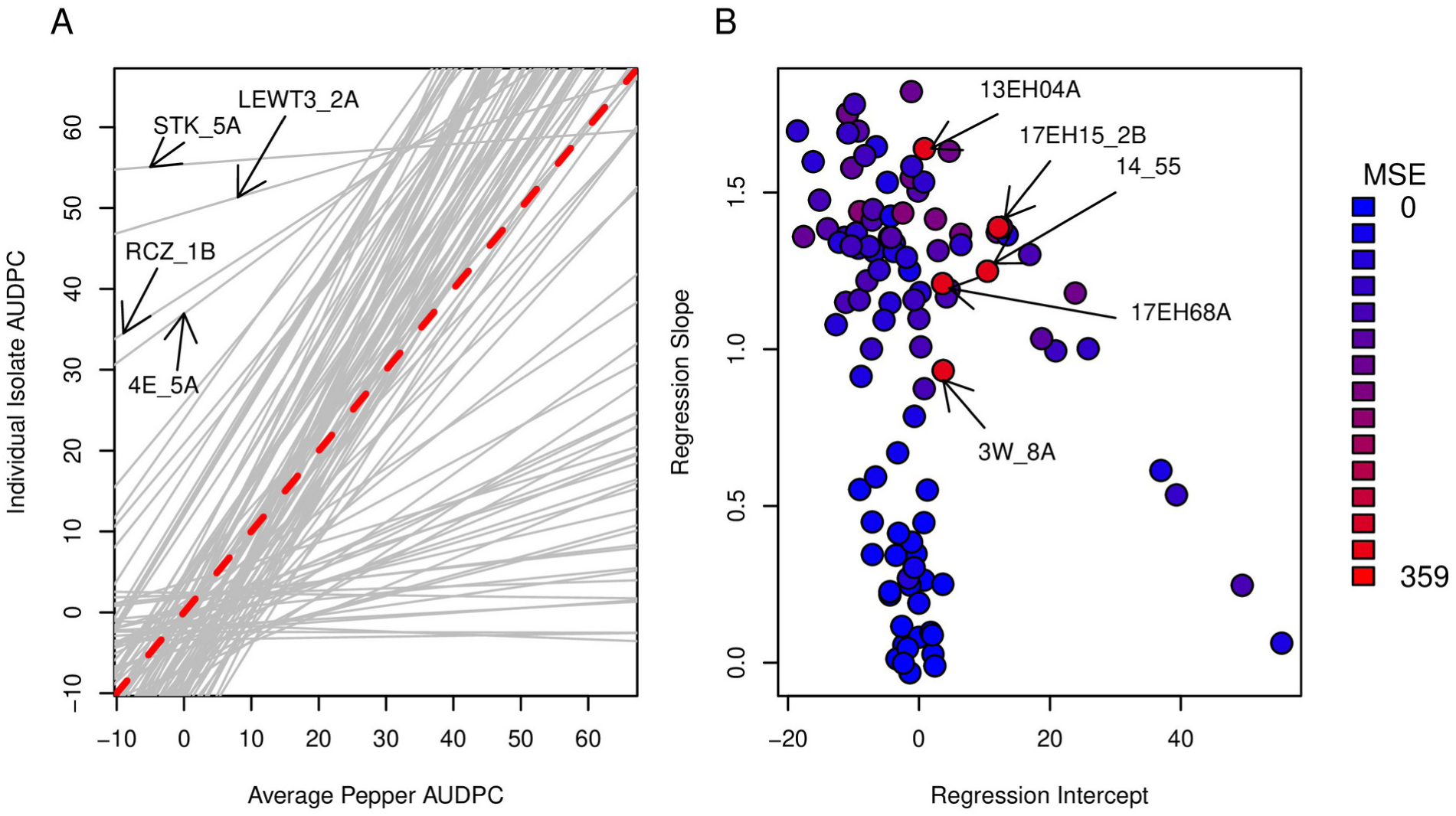
Finlay-Wilkinson regression of individual isolate AUDPCs on pepper mean AUDPCs. A) Finlay-Wilkinson regression line. Red, dashed line has intercept of 0 and slope of 1, and represents a hypothetical isolate with average AUDPC on every pepper. B) Finlay-Wilkinson regression intercepts vs slope, colored by MSE.

The majority of regressions featured low MSE, suggesting that for most isolates, the level of disease caused on a particular pepper accession could be predicted accurately by the average disease resistance of that accession (Figure 3B). However, several MSE outliers (13EH04A, 17EH15_2B, 14_55, 17EH68A, 3W_8A) were noticeable, corresponding to isolates whose virulence levels on particular pepper accessions were poorly predicted by the linear model. For example, 3W_8A caused higher than average AUDPC on both the most susceptible (Red Knight and Aristotle) and resistant (Paladin, Archimedes, and Early Jalapeño) pepper accessions, yet caused lower than average AUDPC on intermediately resistant accessions Revolution, Vanguard, and Perennial. 13EH04A, on the other hand, demonstrated the opposite pattern, causing higher than average AUDPC on intermediately resistant accessions but average AUDPC on the rest.

### Association between pathogen population structure and virulence

Little consistency in virulence profiles was visually apparent among isolates that originated from the same field site (Figure 2). For example, of the four isolates demonstrating high virulence across all eight pepper accessions, three originated from Suffolk County in Long Island, NY. However, these were each sampled (from pumpkin or squash crops) on different farms, each of which also contributed isolates with virulence patterns more typical of the rest of the isolates in this experiment.

To further assess the relationship between the population structure of the isolate panel and their virulence levels on the eight pepper accessions, we compared principal component analysis (PCA) plots of both their genotypic data, using a dataset of 63,475 genome-wide SNP markers typed on the 105 isolates, and phenotypic data, using the matrix of isolate × pepper accession AUDPC LS means. K-means clustering of scores on the first four PCs of the genotype matrix, which collectively accounted for 26.02% of the variance in the genetic data, sorted the isolates into five clusters (Figure 4; Figure S2), three of which exclusively represented single counties in New York [Cluster 2: field sites Ontario #1 and #2 (Ontario County, central NY); Cluster 3: Erie #1, #2, and #4 (Erie County, western NY); Cluster 5: Cayuga #1 (Cayuga County, central NY)] (Table S2).

**Figure 4:**
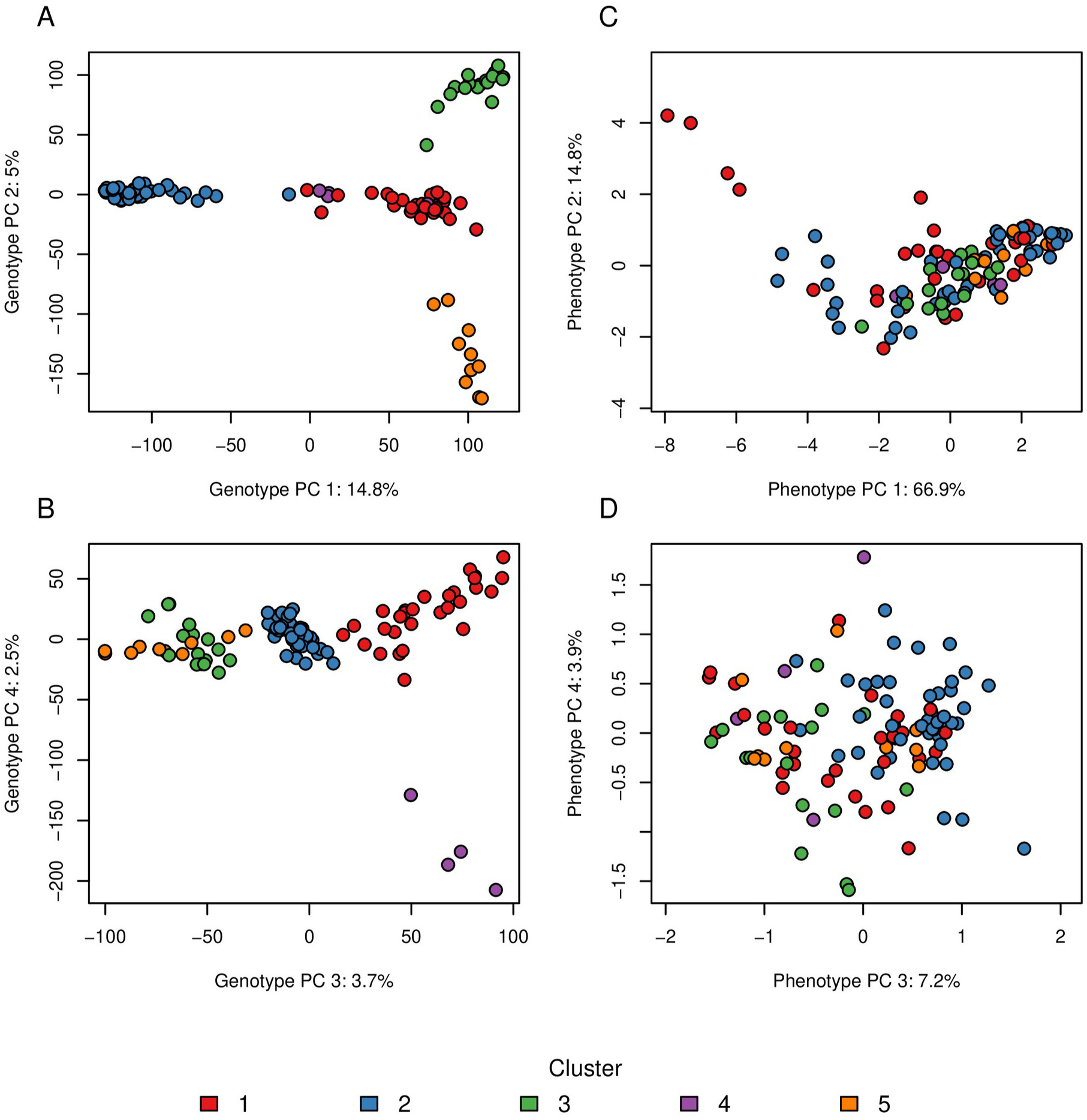
Principal component (PC) analysis plots for A-B) 63,475 single-nucleotide polymorphism markers and C-D) AUDPC estimates on eight pepper accessions, for 105 *P. capsici* isolates. Panes A and C show PC 1 vs PC 2 and panes B and D show PC2 vs PC4 for their respective PCAs. Isolates are colored by genetic cluster, identified by k-means clustering using the four genotype PCs.

Distinct clustering was not evident in the phenotypic PCA plot (Figure 4C-D). Phenotypic PC 1, which explained 66.9% of the variance in the disease data, separated isolates based on their overall virulence, as indicated by its almost perfect correlation with the LS mean for the main effect of isolate (*r* = -0.99). Phenotypic PC 2 explained 14.8% of the variance and appeared to reflect the responsiveness of isolates to changes in average host susceptibility, as suggested by its high correlation with Finlay-Wilkinson slope (*r* = -0.75). Phenotypic PCs 3 and 4 represented combinations of variables that were more difficult to interpret. However, the direction and magnitude of the loadings of individual pepper accessions on these PCs suggested that PC 3 largely measured virulence on Red Knight and Aristotle in relation to virulence on Perennial, Revolution, and Vanguard, whereas PC 4 measured virulence on Perennial and Aristotle in relation to virulence on Revolution and Vanguard (Figure S3).

ANOVAs were conducted to determine if genetic clusters were associated with variation in virulence on individual pepper accessions or on any of the first four phenotypic PCs. Significant differences between genetic clusters were observed for virulence on each of Archimedes, Aristotle, Early Jalapeño, Paladin, and Red Knight, as well as for phenotypic PCs 2, 3, and 4 (Table 3). However, after adjusting for the number of tests following a Bonferroni procedure, differences between genetic clusters were only significant for virulence on Red Knight and for phenotypic PC 3. Of the traits, phenotypic PC 3 showed the strongest association with pathogen population structure, with genetic cluster accounting for 40% of the variance in scores along this component (Table 3). Phenotypic PC 3 mainly differentiated cluster 2 isolates from isolates in clusters 3 and 4 (Figure 4).

**Table 3:** R^2^ values and P-values from models regressing virulence traits on genetic cluster assignment.

| Virulence trait | $R^2$ | $P$-value | Adjusted $P$-value<sup>a</sup> |
| --- | --- | --- | --- |
| Archimedes | 9.73 | $3.8 \times 10^{-2}$ | 0.46 |
| Aristotle | 11.37 | $1.6 \times 10^{-2}$ | 0.19 |
| Early Jalapeño | 9.88 | $3.3 \times 10^{-2}$ | 0.39 |
| Paladin | 13.17 | $6.5 \times 10^{-3}$ | $7.8 \times 10^{-2}$ |
| Perennial | 5.95 | 0.19 | 1 |
| Red Knight | 21.42 | $6.8 \times 10^{-5}$ | $8.2 \times 10^{-4}$ |
| Revolution | 5.78 | 0.21 | 1 |
| Vanguard | 6.41 | 0.16 | 1 |
| PC1 | 8.04 | $7.6 \times 10^{-2}$ | 0.91 |
| PC2 | 10.01 | $3.1 \times 10^{-2}$ | 0.37 |
| PC3 | 40.34 | $1.3 \times 10^{-10}$ | $1.5 \times 10^{-9}$ |
| PC4 | 12.11 | $1.1 \times 10^{-2}$ | 0.13 |

<sup>a</sup> $P$ -values Bonferroni-corrected for multiple tests.

### Genome-wide association studies for host-specific virulence

Genome-wide association studies were conducted for overall virulence averaged across pepper accession, accession-specific virulence on each of the eight accessions retained in the dataset, and phenotypic PCs 2 to 4 (excluding PC 1 due to its high correlation with overall virulence). Because of the varying degree of association between population structure and each virulence trait, a model-testing procedure was used to determine the most appropriate covariates to control for false positive associations due to population structure in each GWAS (Table S3). Traits were log-transformed as needed based on the normality of the residuals from null models fitted with covariates but without individual SNP effects.

Eleven SNPs were identified that surpassed a 10% False Discovery Rate (FDR) threshold for association with one or more virulence traits (Figure 5, Table 4, Table S4). These 11 SNPs were involved in a total of 14 significant marker-trait associations and represented four unique loci on each of scaffolds 1, 37, 39, and 93 of the *P. capsici* reference genome (Lamour et al. 2012a). The loci on scaffolds 1, 37, and 93 contained SNPs that were significantly associated only with overall virulence across pepper accessions, whereas the peak SNP at the scaffold 39 locus was significantly associated with overall virulence in addition to virulence on each of Perennial, Red Knight, and Revolution. While the remaining virulence traits did not result in any significant marker associations (Figure S4), the peak SNPs for the loci on scaffolds 1, 37, 39, and 93 featured in the top 1% of associations for 7, 6, 7, and 4, respectively, of the eight pepper accession-specific virulence traits. In addition, the minor allele at each of the peak SNPs demonstrated consistent effects across pepper accessions; in all four cases, the minor allele was associated with lower virulence on each of the eight accessions (Table S5). Considering only the peak SNPs for each locus, the mean variance explained by each of these four markers for all significant trait associations was 15.06%, ranging from 11.44%, in the case of the peak SNP on scaffold 37 for overall virulence, to 19.12%, in the case of the peak SNP on scaffold 39 for overall virulence (Table S5).

**Figure 5:**
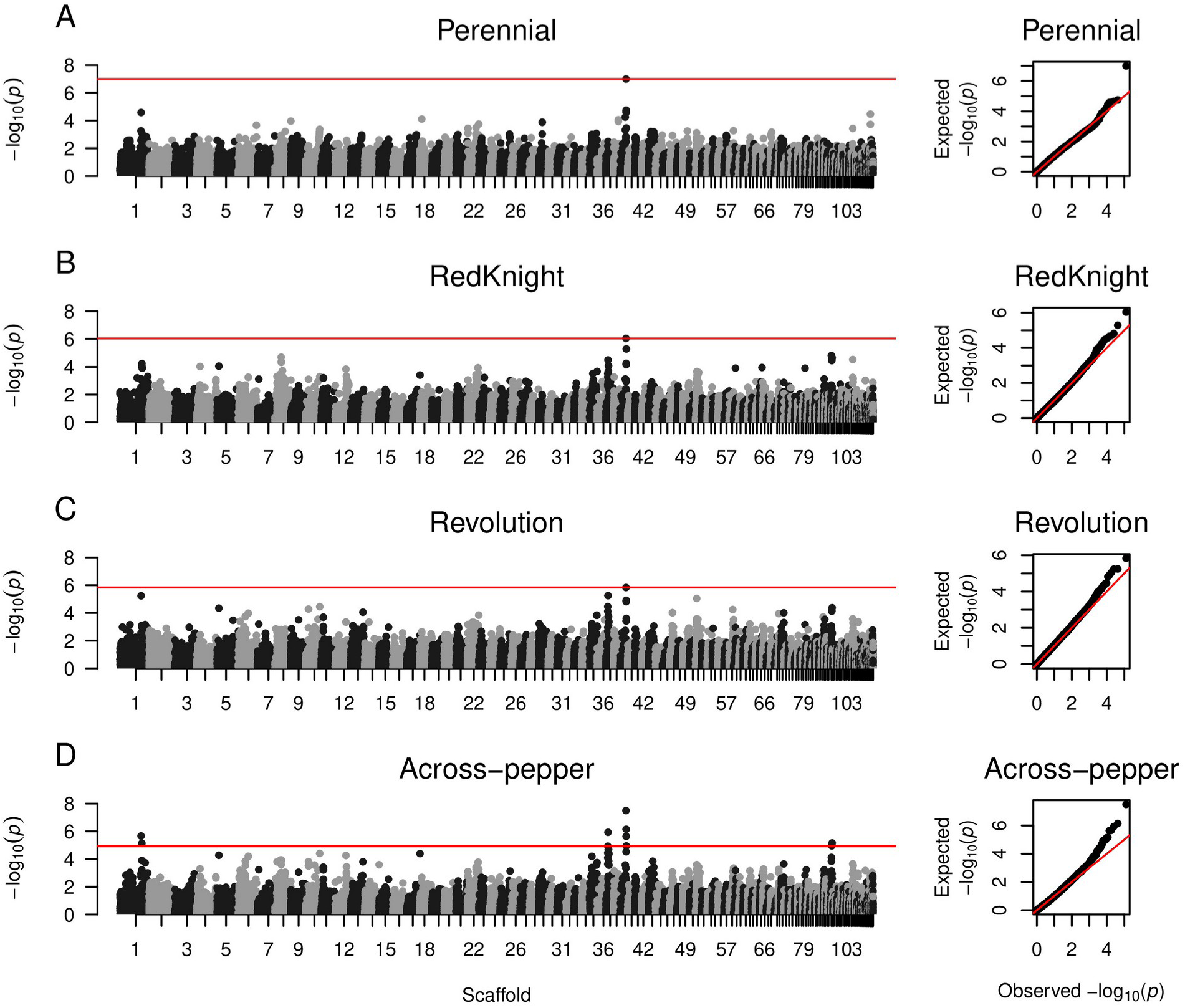
Manhattan and Q-Q plots showing *P*-values from genome-wide association studies for virulence on each of A) Perennial, B) Red Knight and C) Revolution, as well as D) overall, across-pepper virulence.

**Table 4:**
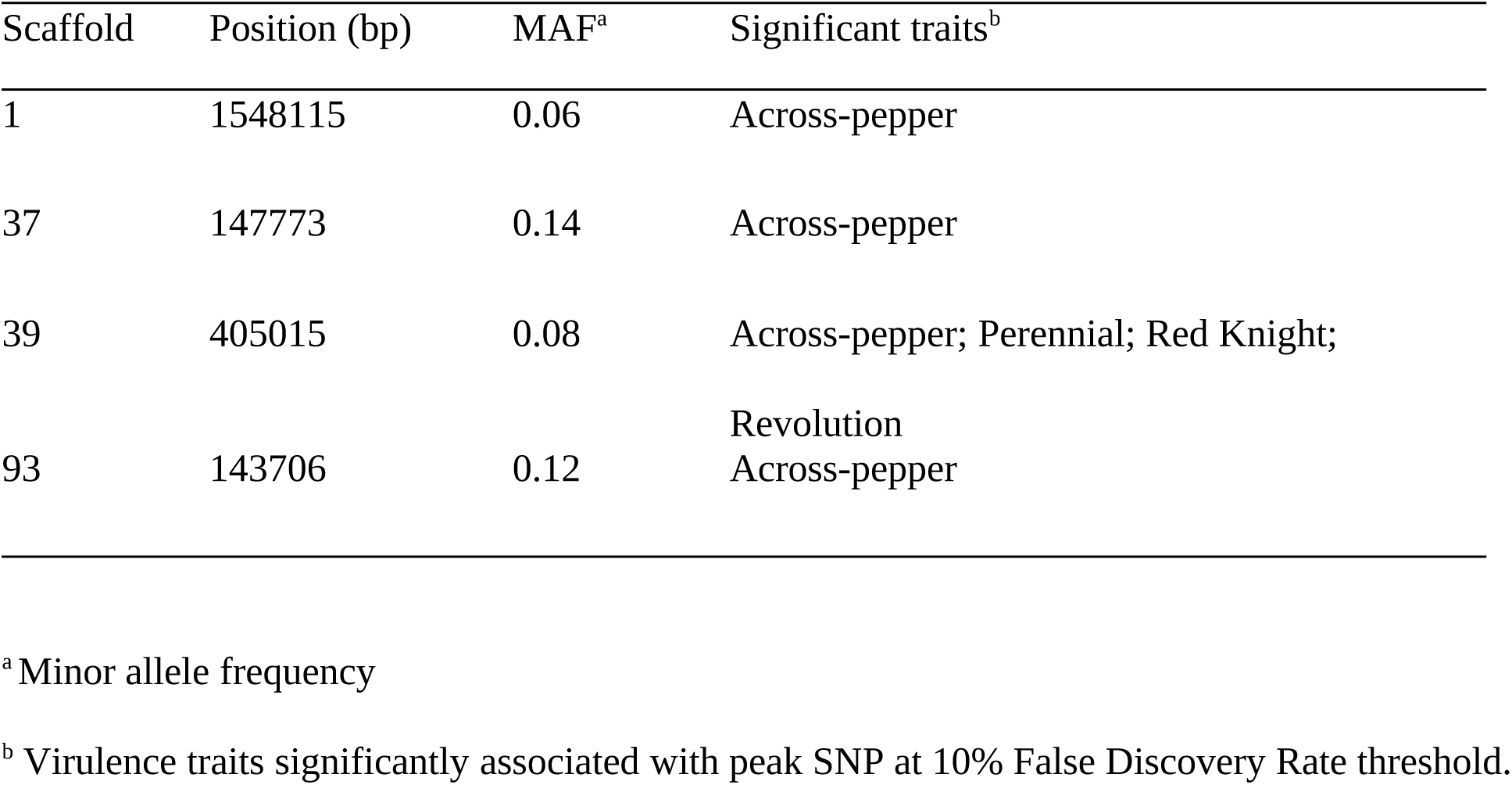
Genomic locations of peak SNPs representing genome-wide association study hits.

Gene annotations within regions in strong linkage disequilibrium (LD) with significant markers were assessed in order to identify potential causal genes underlying the four GWAS signals. In addition to the annotations in the reference genome (Lamour et al. 2012a), we referred to a published list of putative *P. capsici* effectors (Thilliez et al. 2019) and identified their coordinates in the reference genome sequence (Figure 6).

**Figure 6:**
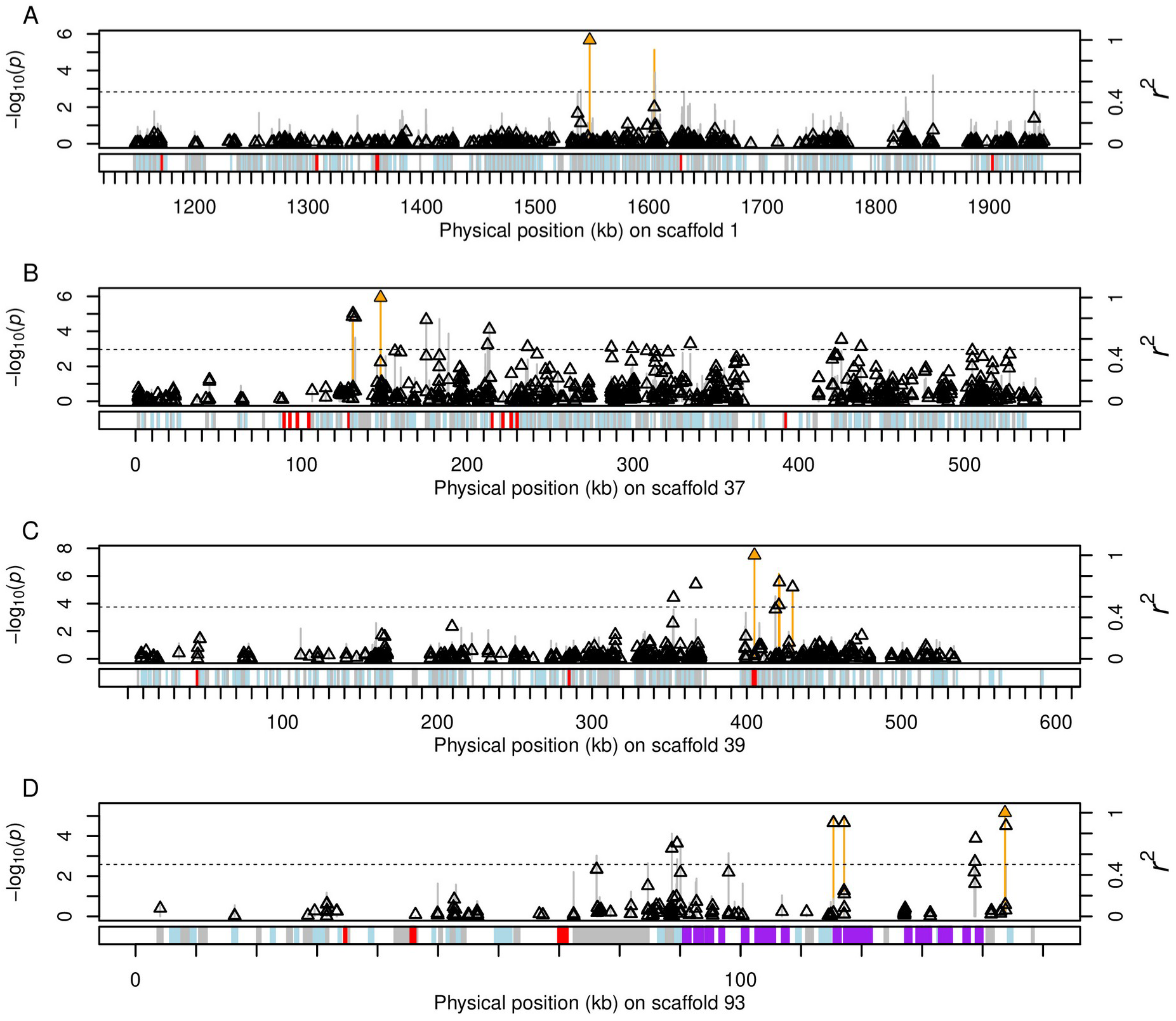
Scatterplots showing genome-wide association results for genomic regions within up to 400 kb from peak SNPs on scaffolds A) 1, B) 37, C) 39, and D) 93 of the *P. capsici* genome. SNP *P*-values for association with across-pepper virulence are represented by vertical lines, with lines for significant SNPs colored in orange. Linkage disequilibrium (*r*^2^) values with the peak SNP on each respective scaffold are represented by triangles, with the triangle for the peak SNP colored in orange. The horizontal dashed line represents *r^2^*=0.5.Tracks below plots show the locations of all annotated genes (Lamour et al. 2012a) alternating in gray and light blue. Putative effectors (Thilliez et al. 2019) are highlighted in red and putative polygalacturonases are highlighted in purple.

The scaffold 1 locus featured two significant SNPs, 1_1548115 and 1_1605065, which were located, respectively, inside a putative Acyl-CoA dehydrogenase (*fgenesh2_kg.PHYCAscaffold_1_#_243_#_Contig762.1*) and a putative transcription initiation factor (*fgenesh2_kg.PHYCAscaffold_1_#_256_#_4100336:1*). These two SNPs were separated by 60 kb and in moderate LD (*r^2^* = 0.35) with each other (Figure 6A). No SNPs in the intervening or surrounding regions were in strong LD (*r^2^* > 0.5) with the peak SNP, and no genes in the immediate area featured annotations with obvious virulence associations.

The scaffold 37 locus, in contrast, was located inside a region of slow LD decay. Only two SNPs, separated by 7 kb, surpassed significance thresholds: 37_131151, inside a gene with a predicted role in protein transport, and 37_147773, which was intergenic. However, SNPs in strong LD (*r^2^* > 0.5) with peak SNP 37_147773 extended over an approximately 300 kb region from 130,776 bp to 437,445 bp (Figure 6B). Ten putative RXLR effector genes were located within 400 kb of peak SNP 37_147773, six of which were within or close to the region spanned by SNPs in strong LD with 37_147773: PcRXLR239, PcRxLR240, PcRxLR241, PcRxLR242, PcRxLR243, and PcRXLR237 (Table S6). PcRXLR240, PcRXLR241, and PcRXLR243 corresponded to reference gene models *estExt2_fgenesh1_pg.C_PHYCAscaffold_370037*, *fgenesh1_pg.PHYCAscaffold_37_#_39,* and *fgenesh1_pg.PHYCAscaffold_37_#_41*, whereas the genomic coordinates for PcRXLR242 and PcRXLR237 overlapped but did not perfectly coincide with reference gene models *fgenesh1_pg.PHYCAscaffold_37_#_40* and

*fgenesh1_pg.PHYCAscaffold_37_#_85,* respectively. PcRXLR239 did not overlap with any gene annotation in the reference genome. Of the five putative RXLRs in the candidate gene search space that corresponded with a reference genome annotation, all but *fgenesh1_pg.PHYCAscaffold_37_#_85* were annotated with a signal peptide.

The peak SNP on scaffold 39, 39_405015, was located inside a gene, *fgenesh1_pg.PHYCAscaffold_39_#_56*, that lacked functional annotation in the *P. capsici* reference genome. However, the location of this gene overlapped with the coordinates for putative RXLR-type effector PcRxLR251 (Figure 6C; Table S6). Both annotations agreed on the same ending position in the reference genome, but the predicted start codon for PcRxLR251 was extended by 348 base pairs compared to the gene model in the reference genome. Although *fgenesh1_pg.PHYCAscaffold_39_#_56* was not annotated with a signal peptide, it appeared to be in a cluster of genes encoding putative secreted proteins, with seven genes annotated with signal peptides within a 40 kb region surrounding the SNP. Of the three other significant SNPs comprising the signal on scaffold 39, two (39_420785 and 39_421156) were located in a putative Trans-2-enoyl-CoA reductase (*e_gw1.39.18.1*) and another (39_429653) was intergenic. Five other SNPs on scaffold 39 were in strong LD (*r^2^* > 0.5) with SNP 39_405015. These SNPs spanned a region of approximately 77 Kb that included 27 additional annotated genes, none of which showed an obvious virulence role.

Three significant markers comprised the scaffold 93 locus, spanning 28 kb from 115,350 bp to 143,706 bp, almost at the end of the scaffold. Seven markers in strong LD (*r^2^* > 0.5) with peak SNP 93_143706 extended over a region of approximately 55 kb, beginning at 89 kb on scaffold 93. Three putative RXLR effectors were located on this scaffold, one of which, PcRxLR496, was located between 70-71 kb, just outside of this region of strong LD. Notably, however, the region of strong LD with 93_143706 co-localized with a cluster of 15 genes all featuring annotations as putative polygalacturonases, which spanned a region of approximately 50 kb from 90 kb to 140 kb (Table S6). One of the three significant SNPs that comprised the GWAS signal, 93_117106, was located inside one of the putative polygalacturonase genes in this cluster (*fgenesh1_pg.PHYCAscaffold_93_#_20*). The other two significant SNPs, 93_115350 and 93_143706, were intergenic, although 93_115350 was located 12 bp upstream from the start codon of another of the putative polygalacturonase genes (*fgenesh1_pg.PHYCAscaffold_93_#_20*).

## DISCUSSION

Genotype-genotype interactions between pepper and *P. capsici* have been of interest to pepper breeders and pathologists for many years (Barchenger et al. 2018a). However, while the genetic control of disease resistance has been studied extensively in pepper (Mallard et al. 2013; Ogundiwin et al. 2005; Rehrig et al. 2014; Siddique et al. 2019; Thabuis et al. 2003; Truong et al. 2012), little is known about the genetic variation in *P. capsici* associated with the ability to cause varying levels of disease on distinct host genotypes. In this study, we measured the disease outcomes for 1,864 distinct combinations of 16 pepper accessions and 117 *P. capsici* isolates. By combining this phenotype data with genotypes at over 60,000 SNP loci in the *P. capsici* isolate panel, we characterized how virulence profiles vary within and between pathogen subpopulations, and identified four loci significantly associated with natural variation for virulence on multiple pepper genotypes.

### Interactions between pepper and *P. capsici*

Of the 16 pepper accessions inoculated in this experiment, eight were largely uninformative for differentiating isolate virulence levels, as they showed complete resistance to almost the entire isolate panel (Figure S1). Surprisingly, among these largely resistant accessions were the six NMRILs that have shown differential reactions to populations of *P. capsici* collected in various locations including New Mexico, Brazil, Taiwan, and Mexico (Barchenger et al. 2018b; da Costa Ribeiro and Bosland 2012; Reyes-Tena et al. 2019; Sy et al. 2008). Only six isolates in our experiment (17EH03_2B, 17EH64A, 3W_2A, AND_4A, LEWT3_2A, and STK_5A) caused any disease on one or more of these NMRILs in both phenotypic replicates. Similarly, Hu et al. (2013) failed to observe any disease on eight NMRILs that they inoculated with 42 *P. capsici* isolates collected in China. These results suggest that avirulence genes that are polymorphic in other geographic populations of *P. capsici* may be largely fixed among the predominantly New York isolates characterized here and the Chinese isolates characterized by Hu et al. (2013). Consequently, different sets of differential hosts are needed to provide relevant information for distinct *P. capsici* populations.

In addition to the NMRILs, several other pepper accessions in this experiment showed unexpected responses to the 117 isolates with which they were challenged. Perennial, for example, which has been used as a resistant parent in mapping populations (Lefebvre and Palloix 1996; Thabuis et al. 2003), featured the third highest median AUDPC (Table 2; Figure 1) of the 16 pepper accessions in this experiment. Early Jalapeño, on the other hand, described by others as susceptible or possessing low levels of resistance to *P. capsici* (Rehrig et al. 2014), performed similarly in our experiment to the bell pepper hybrids Paladin and Archimedes, both of which are marketed as intermediately resistant (Table 2; Figure 1). The high level of resistance of Early Jalapeño to the isolates in our panel also agrees with the unexpectedly consistent resistance levels of the NMRILs, as Early Jalapeño is one of the parents of the NMRIL population in addition to highly resistant landrace CM 334. It is interesting to note that while broad-sense heritabilities for virulence were high on all eight of the pepper accessions retained in our dataset, they were lowest on Early Jalapeño and Perennial (0.72 and 0.79, respectively), meaning that a relatively larger proportion of the variance in disease observed on these peppers could not be attributed to genetic variation among isolates. Of the eight accessions retained for analysis, these two accessions were the only that were not F1 hybrid varieties. One possible explanation for this slight decrease in heritability could be that these two seed sources were partially heterogenous and segregating for disease resistance, as has been observed within open-pollinated accessions of other species (Davis et al. 2007; Grumet and Colle 2017).

Significant genotype-genotype interactions were observed between the eight pepper accessions and 105 *P. capsici* isolates included in our analysis (Table 1). However, 66.9% of the variance in the disease caused by the isolates on the eight accessions (PC 1 of the phenotypic PCA; Figure 4) could be largely explained by the isolate main effect, or the average virulence level of each isolate. An additional 14.8% of the phenotypic variance was strongly associated with the Finlay-Wilkinson regression slope for each isolate, which measures the rate of increase in disease caused by an isolate as the average susceptibility of its host increases. These results indicate that most of the variation in disease observed in our experiment could be predicted by three parameters: the average virulence of isolate, the average resistance of pepper accession, and the rate at which an isolate’s virulence increases with average pepper susceptibility.

Although several isolates featured virulence profiles that showed poor linear relationships with average pepper accession susceptibility (i.e., the outliers in Figure 3b whose MSE values were between 2-4 standard deviations above the mean MSE across isolates), there was little evidence of clear, qualitative crossover interactions, as would be expected in a gene-for-gene system where interactions between R and effector alleles resulted in the complete presence or absence of disease for a given isolate-pepper genotype combination. For example, while we observed several isolates that were able to cause high levels of disease on the mostly resistant bell pepper hybrids Paladin and Archimedes, these isolates were also among the most virulent on the other six pepper accessions. In the classic gene-for-gene model, mutations in an effector gene that enable a pathogen isolate to infect a host with a matching R gene would not also confer increased virulence on separate hosts without that R gene.

However, consistent with the identification in pepper of minor-effect resistance QTL with isolate-specific effects (Ogundiwin et al. 2005; Rehrig et al. 2014; Siddique et al. 2019; Truong et al. 2012), we found evidence in our dataset of smaller, quantitative genotype-genotype interactions. Phenotypic PC 3, for example, appeared to differentiate isolates in terms of their virulence on intermediately resistant pepper accessions Revolution, Vanguard, and Perennial in relation to their virulence on susceptible cultivars Red Knight and Aristotle (Figures 4 and S3). The isolates with the most negative scores along PC 3 (3W_8A, 18051_1A, 17DH16A) caused lower than average disease on the intermediately resistant pepper accessions, but higher than average disease on Red Knight and Aristotle, whereas the isolates with the highest scores (17EH68A, 17EH15_2B, 17EH64A) caused either equivalent levels of disease on the five accessions or even higher disease on Revolution, Vanguard, and Perennial compared to Red Knight and Aristotle (Figure 2). Phenotypic PC 4 differentiated between the intermediately resistant pepper accessions, separating isolates with high virulence on Perennial compared to Vanguard and Revolution (e.g., 568OH and 17EH30B) from those with high virulence on Vanguard and Revolution compared to Perennial (e.g. 17PZ11A and 17PZ02A). Although PCs 3 and 4 collectively explained a relatively small portion of the variance in the data (11.1%), the phenotypic patterns they reflect suggest that the isolates we characterized were polymorphic for virulence loci with minor effects on specific host accessions, in particular Vanguard, Revolution, and Perennial.

### Variation in virulence within and between subpopulations

PCA of the isolate genotype data clearly separated the pathogen population into five genetic clusters that were strongly associated with sampling location in New York (Figures 4A and S2; Table S2). While there was some evidence for differentiation in virulence levels on individual pepper accessions between clusters, the magnitude of phenotypic differences between subpopulations was slight, with genetic cluster explaining 6-21% of the variance in virulence, depending on the accession (Table 3). Phenotypic PC 3, however, showed a much stronger relationship with genetic cluster than did virulence on any individual pepper accession. This indicates that pathogen subpopulations varied in terms of their relative virulence levels on particular combinations of pepper accessions, even if they did not vary in their absolute virulence levels on those accessions. For example, given an isolate from Erie County (genetic cluster 3) and an isolate from Ontario County (genetic cluster 2) with identical virulence on Red Knight, the Ontario County isolate would be more likely to have higher virulence on Vanguard, Revolution, and Perennial (Figure 2). However, without controlling for Red Knight, there was no difference between these counties for absolute virulence levels on Vanguard, Revolution, or Perennial.

Of the virulence traits on individual pepper accessions, the greatest amount of differentiation between genetic clusters was for virulence on Red Knight, the most susceptible accession on average in this experiment. Variation between clusters for their virulence on Red Knight was largely driven by the inclusion of low-virulence isolates in clusters 1, 2, and 4, but not 3 and 4. Within clusters 1, 2, and 4, and even within individual field sites in those clusters, virulence on Red Knight was highly variable. For example, a dramatic example of within-field variation could be observed among the two isolates collected from the Suffolk County #3 site, a pumpkin field sampled in 2018. These two isolates contrasted drastically in their virulence profiles, with LEWT3_2A among the most virulent isolates on all eight pepper accessions and LEWT3_6A either avirulent or lowly virulent on the eight accessions (Figure 2). It is unclear what evolutionary forces would promote the maintenance of high variation for virulence within single, isolated subpopulations, although a similar phenomenon has been observed in fungal pathogen *Zymoseptoria tritici* (Dutta et al. 2021). One possibility is that diversifying selection acts to favor both highly virulent isolates with increased fitness on healthy hosts as well as less virulent, more saprophytic isolates that are preferentially able to colonize dead tissue, as shown in fungal barley pathogen *Rynchosporium secalis* (Abang et al. 2006).

It is also possible, considering the wide host range of *P. capsici* and the diverse crops grown on most vegetable farms where it is found in New York, that there are tradeoffs in virulence on different host species, with certain genetic variants having opposing effects on fitness depending on the plant host. In this scenario, selection would favor different sets of alleles depending on the crop planted to a field in a given year, but fluctuating selection due to crop rotation as well as the presence of a bank of oospores in the soil, which germinate asynchronously and have been shown to act as a reservoir of genetic diversity (Carlson et al. 2017), would maintain genetic variation in a population for virulence on different crops. It is interesting to note that most of the isolates in our study were sampled from squash or pumpkin, with Erie County #2 the only field site where isolates were sampled from a pepper crop. Erie County #2 was also one of only two sites (excluding sites with <3 isolates) where every isolate was observed to cause high disease on Red Knight, consistent with the hypothesis that within-year selection acts to favor isolates virulent on that particular host crop. In contrast, isolates collected in 2017 from Ontario #1, a site that experienced two consecutive disease epidemics on pumpkin crops and had reportedly never been planted to pepper since inoculum was introduced to the site, were divided almost 50:50 in terms of those with high and low virulence on Red Knight. However, evidence elsewhere of host species specialization in *P. capsici* is inconclusive. Although some studies have suggested a relationship in *P. capsici* between host of origin and virulence on a particular host species (Lee et al. 2001; Ristaino 1990), other studies have found no evidence of a link (Enzenbacher and Hausbeck 2012; Yin et al. 2012). Segregation ratios in inheritance studies conducted by Polach and Webster (1972) indicated the existence of genes controlling pathogenicity on different hosts, but little further research has been conducted on the genetic control of host species specialization in *P. capsici*.

### Genome-wide association study results

We identified four loci that were significantly associated with overall virulence, one of which was also significantly associated with virulence on each of Perennial, Red Knight, and Revolution (Figure 5,6; Table 4). For three of these loci - all but the region on scaffold 1 - candidate genes with a potential virulence function were identified within the area of LD decay with the peak SNP (Table S6).

Six putative RXLR effectors (PcRXLR237, PcRXLR239, PcRXLR240, PcRXLR241, PcRXLR242, and PcRXLR243) were identified as candidates for the causal gene underlying the virulence locus identified on scaffold 37. Expression of five of these six genes (all but PcRXLR243) was detected during infection of tomato (Jupe et al. 2013), and all but PcRXLR239 and PcRXLR237 were found to be expressed to varying amounts in different *P. capsici* structures *in vitro* (Chen et al. 2013). BLAST searches revealed that PcRXLR239, PcRXLR240, and PcRXLR243 shared high homology with genes in other *Phytophthora* species, whereas PcRXLR237, PcRXLR241 and PcRXLR242 did not. In the case of the scaffold 37 locus, the peak SNP (39_405015) was actually located inside an RXLR effector, PcRXLR251, corresponding to gene model *fgenesh1_pg.PHYCAscaffold_39_#_56* in the *P. capsici* reference genome (Lamour et al. 2012a). Expression of this gene was detected *in vitro* in low amounts in mycelia, zoospores, and germinating cysts (Chen et al. 2013), although it was not detected during infection of tomato (Jupe et al. 2013). A BLAST search showed that this gene featured homologs in the genomes of several other *Phytophthora* spp.

Unlike the scaffold 37 and 39 loci, no putative RXLR genes were located within the virulence-associated region mapped on scaffold 93. However, significant SNPs in this locus were located amid a cluster of 15 putative polygalacturonase genes. Polygalacturonases are types of cell wall degrading enzymes that macerate plant tissue through the hydrolysis of polygalacturonic acid, a major component of pectin in cell walls. The secretion of polygalacturonases has been shown to enhance the virulence of many plant pathogenic bacteria, fungi, and oomycetes, including that of *Phytophthora capsici* (Lei et al. 1985; Li et al. 2012; Shieh et al. 1997; Sun et al. 2009). Polygalacturonases trigger host defense responses indirectly via plant recognition of their hydrolysis products (Pilling and Hofte 2003) and are also directly targeted by plant polygalacturonase-inhibiting proteins that bind them and inhibit their activity (Di Matteo et al. 2006).

In the case of all four loci discovered, the minor allele at each peak SNP, ranging in frequency from 0.06 on scaffold 1 to 0.14 on scaffold 37, was associated with lower virulence. The reduction in virulence caused by the causal mutation or mutations at each of these loci could be mediated by a reduction in the activity of their respective gene products, or alternatively, by their increased detection by host plant defenses. For example, assuming the causal gene at the scaffold 93 locus is a polygalacturonase, the low-virulence allele may contain a mutation in its active site impacting its ability to hydrolyze polygalacturonic acid, or alternatively, may contain a mutation that allows it to be more easily bound and inhibited by polygalacturonase-inhibiting proteins. Similarly, a mutation in an RXLR effector may either directly impact its virulence function or enable its recognition by a cognate R gene in the plant host.

Each of the four loci that we mapped was associated with broad-spectrum virulence, in that their minor alleles showed a directionally consistent impact on disease outcomes on all eight pepper accessions retained in our analysis, including on the highly susceptible cultivar Red Knight. The universally negative effect of these alleles on virulence suggests that selection would act strongly against them. However, it is possible that these alleles may confer a fitness advantage on other host species or even other pepper genotypes not included in this experiment. Functional characterization of the candidate genes underlying these loci is necessary to understand how they are associated with variation in virulence, and what, if any, fitness advantages are conferred by the allele conferring low virulence on pepper.

The presence of significant genotype-by-genotype interactions indicates that there must be loci in *P. capsici* with differential effects on virulence for different pepper hosts. It is possible that our experiment lacked sufficient statistical power to detect these accession-specific virulence loci. Our sample size of 105 individuals is low for GWAS, although similar sample sizes have been used successfully for mapping quantitative virulence loci in fungal species *Zymoseptoria tritici* (Hartmann et al. 2017). Other studies have reported highly polygenic architectures for virulence in several fungal species (Hartmann et al. 2017; Soltis et al. 2019). The results from our study support a model for *P. capsici* in which virulence loci of medium to large effect act independently of host genotype, whereas host genotype-specific virulence loci are of smaller effect and consequently more challenging to identify via GWAS. This would imply that the genetic architecture of pathogen virulence is similar to the genetic architecture of resistance in the host, as loci in pepper with large effects on *P. capsici* resistance are broad-spectrum but smaller-effect loci have isolate-specific effects (Mallard et al. 2013; Ogundiwin et al. 2005; Rehrig et al. 2014; Siddique et al. 2019; Truong et al. 2012). Alternatively, allelic heterogeneity may be another possible explanation for the lack of accession-specific associations detected in this experiment. In several fungal pathogens, high mutation rates, large population sizes, and positive selection have led to allelic diversification at avirulence loci, with multiple independent mutations in the same genes conferring the same phenotypic effect (Dai et al. 2010; Daverdin et al. 2012). This scenario would lead to multiple, rare causal variants, each in weak LD with tested SNP markers and therefore difficult to detect with all but impractical sample sizes.

### Conclusions

We identified significant evidence for genotype-genotype interactions in the pepper-*P. capsici* pathosystem. However, the magnitude of these interactions were quantitative rather than qualitative, and most of the variation in the virulence profiles of 105 isolates could be explained by their average virulence across host genotypes. Consistent with the quantitative nature of these interactions, we were unable to detect any SNPs significantly associated with differential virulence, suggestive of many genes of small effect playing a role in host specialization. However, we were able to map four regions of the *P. capsici* genome implicated in variation in broad-spectrum virulence on multiple pepper accessions. Three of these four loci colocalized with genes whose annotations provided biological support to their virulence associations, whether as RXLR effectors in the case of the loci on scaffolds 37 and 39 or cell wall degrading polygalacturonases in the case of the scaffold 93 locus. It is unclear why variation at these loci would be preserved in *P. capsici*, although we speculate that the alleles conferring lower virulence on pepper may confer a fitness advantage on different host species or in a different stage of the *P. capsici* life cycle. The results of this project are expected to inform pepper resistance breeding and the deployment of resistant varieties, in addition to providing a foundation for the functional validation of genes associated with virulence variation in *P. capsici*.

## MATERIALS AND METHODS

### Pepper accessions

Seed of commercial varieties included in this experiment were obtained directly from seed companies [Early Jalapeño: Johnny’s Selected Seeds (Winslow, ME); Archimedes, Aristotle, and Red Knight: Stokes Seeds (Thorold, Ontario, CA); Paladin: Syngenta (Greensboro, NC); Intruder, Revolution, and Vanguard: Harris Seeds (Rochester, NY)]. Small quantities of seed of the remaining accessions were obtained from several sources and increased in a greenhouse at Cornell Agri-Tech in Geneva, NY, in 2019. Seed for the six NMRIL lines were obtained from Dr. Paul Bosland, seed for CM334 from Dr. Michael Mazourek, and seed for Perennial (PI 631147) from the U.S. National Plant Germplasm System. Three to four plants of each accession were grown in 3-gal pots and flowers were vibrated daily in order to promote self-pollination. Mature fruit from all plants within each accession were then bulked and seed was extracted, washed with 10% trisodium phosphate, and dried at 24-28 °C for 1-2 days.

The six NMRIL lines included here were selected from the complete set of 76 NMRILs by consulting previously published data from race characterization experiments (Barchenger et al. 2018b; Monroy-Barbosa and Bosland et al. 2011; Sy et al. 2008) and identifying lines whose resistance responses had high variances and low correlations between each other. For the first replicate of the experiment, only three bell pepper hybrids - Red Knight, Aristotle, and Paladin - were included. These varieties were chosen for their known, varying levels of overall Phytophthora root and crown rot resistance, with Red Knight highly susceptible and Aristotle and Paladin possessing low and intermediate levels of resistance, respectively (Dunn et al. 2013, 2014; Krasnow et al. 2017; Parada-Rojas and Quesada-Ocampo 2019). Because four of the NMRIL lines (NMRIL-A, NMRIL-H, NMRIL-I, and NMRIL-Z) showed complete resistance to almost all of the isolates in the first replicate of this experiment, they were replaced in the second replicate with four additional bell pepper hybrids listed as intermediately resistant or tolerant to Phytophthora root rot in seed catalogs (Archimedes, Intruder, Revolution, and Vanguard).

### Pathogen isolates

The 129 genetically unique *P. capsici* isolates described in Vogel et al. (2021) were assayed for their degree of zoospore production and a subset of 117 isolates was identified that consistently sporulated in culture (data not shown). Two of these 117 isolates, 14-55 and 17PZ21A, were not included in the clone-corrected set described in Vogel et al. (2021), but were indirectly represented by a single-zoospore progeny in the case of 14_55 (14_55C) and by another isolate of the same clonal lineage in the case of 17PZ21A (17PZ18A). Isolates were transferred from long-term hemp seed storage tubes (Vogel et al. 2021) to PARPH plates (Jeffers and Martin 1986) prior to conducting each replicate of disease assays. These PARPH plates were wrapped with parafilm (Beemis, Neenah, WI) and stored at room temperature until used for transferring of plugs for inoculum preparation.

### Experimental design

Disease assays were conducted in greenhouses at Cornell Agri-Tech that were maintained at 29 °C day/ 23 °C night. The experiment was laid out as a split-plot design with pepper accession as the sub-plot treatment and pathogen isolate as the whole-plot treatment. Whole plots consisted of single 72-cell trays, with experimental units consisting of six-plant plots of peppers. Each tray, filled with soilless potting media, contained twelve plots randomly assigned one of the twelve pepper accessions included in each rep. Each tray was inoculated with one pathogen isolate.

Because of space and inoculum production constraints, the experiment was conducted in five batches, or incomplete blocks, per replicate. Each block contained 22-26 trays each inoculated with one of the 117 experimental isolates, in addition to one tray inoculated with water as a negative control, and three trays each inoculated with one of three check isolates. The three check isolates consisted of two isolates also represented among the 117 experimental isolates (SJV_CAA and 17EH01C) and an additional isolate (0664-1; Dunn et al. 2010). These checks were chosen for their differing degrees of overall virulence, based on preliminary data, and were included in every block in order to measure block-to-block variation in disease severity. Two complete replicates of the experiment were conducted. However, as mentioned previously, four of the 12 pepper accessions in Rep 1 were replaced with a different set of four accessions in Rep 2, resulting in eight pepper accessions with two observations for each isolate and eight pepper accessions with only one observation for each isolate. In addition, three isolates failed to sporulate in one of two replicatess and were therefore only included in one replicate.

Pepper plants were inoculated at five weeks of age. Prior to inoculation, any cells of a tray where seed did not germinate were replaced with plants from a back-up set in order to ensure that all plots contained six plants. Plants were inoculated with a zoospore suspension that was pipetted to the potting soil surface directly adjacent to each pepper stem. Each plant was inoculated with a total of 10^5^ zoospores. To prepare inoculum, isolates were plated on 15% unfiltered V8 agar and incubated at room temperature with 15 h of fluorescent lighting per day for 7, 10, or 14 days, depending on the optimal incubation period to maximize sporulation for each isolate. Plates were flooded with distilled water and an L-shaped spreading rod was used to dislodge sporangia from the surface of plates, which were then collected in flasks and incubated at room temperature for 30-60 min to promote the release of zoospores. Zoospore concentrations were measured using a hemocytometer and solutions were diluted to the desired final concentration for inoculating.

Plots were rated for incidence of mortality at 4, 6, 8, 11, 13, and 15 days post inoculation (dpi). Plants were declared dead when they had fewer than two non-wilting, fully expanded leaves attached. Mortality ratings at the six time points were then used to calculate the AUDPC (Shaner and Finney 1977) for each plot.

### Statistical analysis of virulence experiment

The dataset was filtered to remove observations corresponding to negative controls, isolates that did not cause disease on any pepper accession, or pepper accessions on which fewer than 20% of isolates were able to kill at least one plant in both reps. The following mixed linear model was then fit:

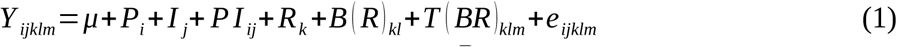

where *Y _ijklm_* is an individual observation, *μ* is the intercept, *P_i_* is the fixed effect of the *i*th pepper accession, *I _j_* is the fixed effect of the *j*th isolate (including check isolates), *PI_ij_*is the fixed effect of the interaction of the *i*th pepper accession and the *j*th isolate, *R_k_* is the fixed effect of the *k*th replicate, 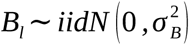 is the random effect of the *l*th incomplete block nested in the *k*th replicate, 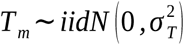 is the random effect of the *m*th tray nested in the l*th* block in the *k*th replicate (e.g. whole plot error term), and 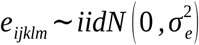 is the residual error effect. This model was fit twice using different subsets of the data. First, in order to conduct significance tests of model terms, it was fit using a balanced subset of the data that featured only pepper accessions included in both replicates. Fixed effects were tested with incremental F tests using the Kenward-Roger approximation for calculation of denominator degrees of freedom (Kenward and Roger 1997). Random terms were tested using likelihood ratio tests. The model was then fit with the full dataset in order to calculate least squares means (LS means) for each isolate, pepper accession, and isolate × pepper accession combination, including for peppers on which only one replicate of each isolate was observed. These LS means were used in all downstream data analyses, such as GWAS.

In addition, the data were sub-divided into separate datasets for each of the pepper accessions included in both reps, in order to calculate broad-sense heritability (*H^2^)* for virulence on these peppers. The following model was fit for each pepper-specific dataset:

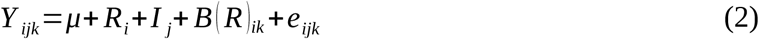

where *Y _ijk_* is an individual observation, *μ* is the intercept, *R_i_* is the fixed effect of the *i*th replicate, 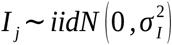 is the random effect of the *j*th isolate, 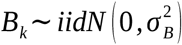 is the random effect of the *k*th incomplete block nested in the *i*th replicate, and 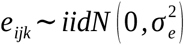 is the residual error effect. Broad-sense heritabilities, calculated on an entry-mean basis, were then estimated using the following formula:

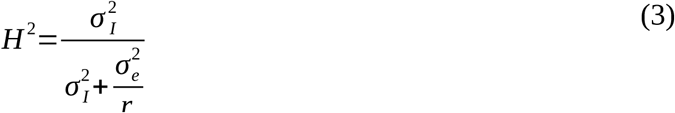

where *r* is the harmonic mean of the number of observations per isolate on a given pepper.

Models were fit using the ASReml-R v 4 package (Butler et al. 2009). LS means were calculated using the *predict()* function. Heritabilities for virulence and their associated standard errors were calculated using the *vpredict()* function.

Virulence stability, or the relationship between the disease caused by individual isolates on a pepper and the average disease severity of that pepper, was assessed by using the following model, originally conceived for the analysis of genotype × environment interactions in crop varieties (Eberhart and Russell 1966; Finlay and Wilkinson 1963):

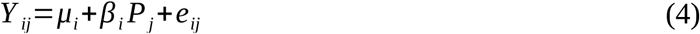

where *Y _ij_* is the LS mean for AUDPC for the *i*th isolate on the *j*th pepper, *μ_i_* is an intercept for the *i*th isolate, *β_i_*is the regression coefficient (i.e. slope) for the *i*th isolate denoting its response to an increase in the overall susceptibility of its pepper host, *P_j_* is the AUDPC LS mean for the *j*th pepper, averaged across isolates, and *e_ij_* is the residual error associated with the *i*th isolate and the *j*th pepper. Three parameters from this model were used to interpret the extent of virulence stability for each isolate: *μ_i_*, the intercept, and *β_i_*, the slope, as in Finlay and Wilkinson (1963), as well as the MSE, as in Eberhart and Russell (1966). These models were fit using the lm() function in R.

### Genotype data

The clone-corrected GBS SNP dataset from Vogel et al. (2021) was subset to include only those isolates that were phenotyped in this experiment and pathogenic on at least one pepper accession. This SNP set was then filtered to remove sites with minor allele frequency < 0.05. Genotypes for isolates 14_55C and 17PZ18A were assigned to the phenotypic records for isolates 14_55 and 17PZ21A, respectively. For all downstream analyses that required complete genotype data, such as GWAS and PCA, missing genotype calls were imputed with the mean allele dosage for that marker.

### Association between population structure and virulence

PCA of both genotypic and phenotypic data (i.e., LS means for isolate × pepper interactions) was performed using the R function *prcomp()*, using centered and unit variance-scaled variables with mean-imputed missing data. The number of genotypic PCs selected for inclusion in analysis was determined by visually identifying the “elbow” in a PCA scree plot. Similarly, to choose the optimal number of clusters for k-means clustering of the genotypic PCA, the k-means algorithm was run with values for the number of clusters ranging from 2 to 10 and the “elbow” was identified in a plot of total within-cluster sum of squares as a function of the number of clusters. ANOVA, using the R functions *lm()* and *anova()*, was conducted to test the association between genetic cluster assignment and each phenotypic PC and virulence trait. Both raw *P*-values and adjusted *P*-values with a Bonferroni correction for the number of ANOVAs performed are reported.

### Genome-wide association study

For each trait used as a response in GWAS, a model-fitting procedure was performed to identify the most appropriate covariates to include to control for population structure, as well as to determine if a log transformation was needed based on analysis of residuals. First, a null model (i.e. a model that did not test individual SNP effects) was fit for each phenotype with up to the first four PCs of the marker matrix as fixed effects and both with and without a random effect for isolate with a covariance structure defined by a genomic relationship matrix (GRM). The GRM was estimated using the *A.mat()* function in the R package rrBLUP (Endelman 2011) using the same SNP set used for GWAS. If the residuals from the best-fitting model, as determined based on the Akaike information criterion (Sakamoto et al. 1986), failed a Shapiro-Wilk test for normality, as implemented in the R function *shapiro.test()*, the phenotype was then log transformed, after first adding a constant to the phenotype to make the minimum value equal to 1. This log transformation was only retained if the model residuals increased in normality, as reflected by the Shapiro-Wilk statistic. Finally, for traits in which a log transformation was elected, the null model-fitting procedure was performed again, in case a different set of covariates demonstrated a better fit with the log-transformed phenotype.

Association tests for individual SNPs were then performed with the identified best-fitting null model for each trait. Markers were declared significantly associated with a trait if their *P*-value surpassed a 10% FDR threshold. Model testing and GWAS were performed using the GENESIS R package (Gogarten et al. 2019). The R package qqman was used to create Manhattan and Q-Q plots (Turner 2014). Allelic effects and R^2^ values for significant markers were calculated from simplified models without any covariates, in which non-transformed phenotypes were regressed on SNP genotypes for that marker.

### Candidate gene identification

Pairwise LD within the population of isolates used in this experiment was previously found to decay to background levels within a distance of approximately 400 kb (Vogel et al. 2021). However, due to variable rates of LD decay across the genome, the LD between the peak SNP at each GWAS signal and all neighboring SNPs within 400 kb was estimated using VCFtools version 0.1.16 (Danecek et al. 2011). Genes with virulence-related annotations were considered as strong candidates at each GWAS signal if they resided within a region spanned by SNPs in strong LD, which we considered as *r^2^* > 0.5, with the peak SNP at that locus.

Gene ontology, KEGG, KOG, signal peptide, and Interpro annotations for gene models from the *P. capsici* reference genome (Lamour et al. 2012a) were assessed in order to identify candidate genes with potential virulence functions. In addition, we referred to a separate list of 573 putative RXLR or CRN effectors used in an effector target enrichment sequencing project (Thilliez et al. 2019). The genomic coordinates of these genes were determined by aligning their sequences against the *P. capsici* reference genome (Lamour et al. 2012) using blastn (Camacho et al. 2009) and identifying the best hit for each gene. In all cases, the best hit featured 100% identity with the queried sequence. Gene IDs for the putative effectors are reported here as in Thilliez et al. (2019).

### Data availability

VCF files with isolate genotype calls and LS means for isolate-pepper accession combinations are publicly available at CyVerse (https://data.cyverse.org/dav-anon/iplant/projects/GoreLab/dataFromPubs/ Vogel_PcapPepper2022). All scripts used in data analysis can be found on Github at https://github.com/gmv23/Pepper-by-Pcap-interactions.

## ACKNOWLEDGEMENTS

Colin Day, Taylere Hermann, Nicholas King, Holly Lange and Carolina Puentes Silva all provided valuable technical assistance for this project. We would also like to thank Drs. Christopher Hernandez and Ryokei Tanaka for their support with statistical analysis. Seed of the non-commercial pepper accessions used in this experiment was kindly provided by Dr. Paul Bosland, Dr. Michael Mazourek, and the USDA ARS Plant Genetics Resources Conservation Unit in Griffin, GA. Funding for this project was provided in part by a grant from the New York State Department of Agriculture and Markets.

## SUPPLEMENTARY TABLES

**Table S1:**
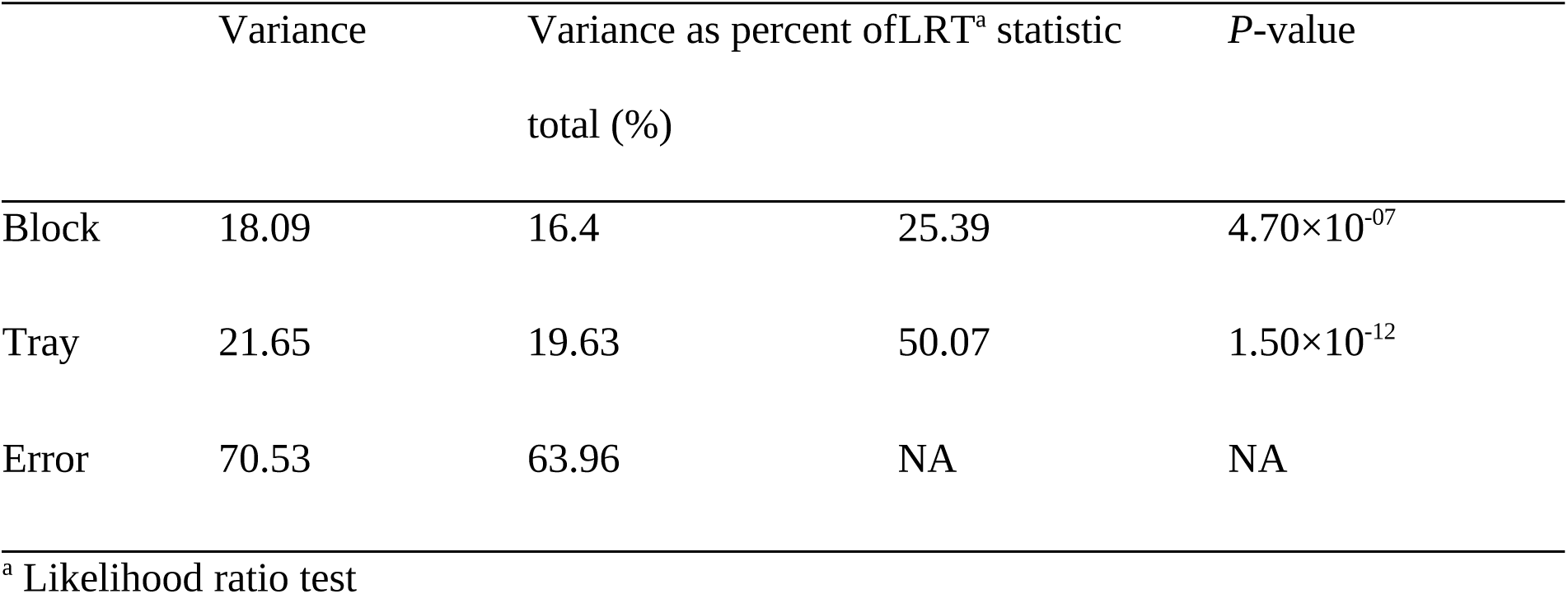
Variances explained by random effects and their test statistics and P-values in model testing the effects of isolate, pepper, and their interaction on AUDPC.

**Table S2:** Number of isolates from each field site or state of origin assigned to each of the five =South Carolina; Sc=Schenectady County, NY; Su=Suffolk County, NY; Ti=Tioga County, NY; To=Tompkins County, NY.

| Field site or state of origin<sup>a</sup> | Genetic cluster |  |  |  |  |
| --- | --- | --- | --- | --- | --- |
|  | 1 | 2 | 3 | 4 | 5 |
| CA | 2 | 0 | 0 | 0 | 0 |
| Ca #1 | 0 | 0 | 0 | 0 | 10 |
| Co #1 | 0 | 0 | 0 | 1 | 0 |
| Er #1 | 1 | 0 | 7 | 0 | 0 |
| Er #2 | 0 | 0 | 9 | 0 | 0 |
| Er #3 | 0 | 0 | 1 | 0 | 0 |
| FL | 1 | 0 | 0 | 0 | 0 |
| Mo #1 | 0 | 0 | 0 | 1 | 0 |
| NM | 1 | 0 | 0 | 0 | 0 |
| NY unknown | 1 | 0 | 0 | 2 | 0 |
| OH | 1 | 0 | 0 | 0 | 0 |
| On #1 | 0 | 43 | 0 | 0 | 0 |
| On #2 | 0 | 2 | 0 | 0 | 0 |
| SC | 4 | 0 | 0 | 0 | 0 |
| Sc #1 | 1 | 0 | 0 | 0 | 0 |
| Sc #2 | 1 | 0 | 0 | 0 | 0 |
| Su #1 | 6 | 0 | 0 | 0 | 0 |
| Su #2 | 2 | 0 | 0 | 0 | 0 |
| Su #3 | 2 | 0 | 0 | 0 | 0 |
| Su #4 | 3 | 0 | 0 | 0 | 0 |
| Su #5 | 1 | 0 | 0 | 0 | 0 |
| Ti #1 | 1 | 0 | 0 | 0 | 0 |
| To #1 | 1 | 0 | 0 | 0 | 0 |

<sup>a</sup> CA=California; Ca=Cayuga County, NY; Co=Columbia County, NY; Er=Erie County, NY; FL=Florida; Mo=Monroe County, NY; NM=New Mexico; OH=Ohio; On=Ontario County, NY; SC=South Carolina; Sc=Schenectady County, NY; Su=Suffolk County, NY; Ti=Tioga County, NY; To=Tompkins County, NY.

**Table S3:** Transformations and covariates used in GWAS models.

| Trait | Log-transformed | GRM/Covariates |
| --- | --- | --- |
| Archimedes | TRUE | None |
| Aristotle | FALSE | PC1+PC2+PC3 |
| EarlyJalapeno | TRUE | PC1+PC2+PC3 |
| Paladin | TRUE | PC1+PC2+PC3+PC4 |
| Perennial | TRUE | None |
| RedKnight | FALSE | PC1+PC2 |
| Revolution | TRUE | PC1+PC2 |
| Vanguard | TRUE | PC1+PC2 |
| Across-pepper | FALSE | PC1+PC2+PC3 |
| PhenoPC2 | TRUE | K |
| PhenoPC3 | FALSE | K+PC1 |
| PhenoPC4 | FALSE | PC1 |

**Table S4:** Significant marker-trait associations.

| Scaffold | Position (bp) | Trait | *P*-value | FDR-adjusted *P*-value |
| --- | --- | --- | --- | --- |
| 1 | 1548115 | Across-pepper | $2.2 \times 10^{-6}$ | 0.03 |
| 1 | 1605065 | Across-pepper | $7.3 \times 10^{-6}$ | 0.07 |
| 37 | 131151 | Across-pepper | $1.2 \times 10^{-5}$ | 0.07 |
| 37 | 147773 | Across-pepper | $1.2 \times 10^{-6}$ | 0.03 |
| 39 | 405015 | Perennial | $1 \times 10^{-7}$ | 0.01 |
| 39 | 405015 | RedKnight | $9.1 \times 10^{-7}$ | 0.06 |
| 39 | 405015 | Revolution | $1.5 \times 10^{-6}$ | 0.09 |
| 39 | 405015 | Across-pepper | $3.2 \times 10^{-8}$ | 0 |
| 39 | 420785 | Across-pepper | $7.2 \times 10^{-7}$ | 0.02 |
| 39 | 421156 | Across-pepper | $2.3 \times 10^{-6}$ | 0.03 |
| 39 | 429653 | Across-pepper | $1.1 \times 10^{-5}$ | 0.07 |
| 93 | 115350 | Across-pepper | $1.1 \times 10^{-5}$ | 0.07 |
| 93 | 117106 | Across-pepper | $1 \times 10^{-5}$ | 0.07 |
| 93 | 143706 | Across-pepper | $6.9 \times 10^{-6}$ | 0.07 |

**Table S5:** R^2^ values and allelic effect of the minor allele for each peak SNP on each virulence trait or phenotypic PC^a^.

|  | 1_1548115 |  | 37_147773 |  | 39_405015 |  | 93_143706 |  |
| --- | --- | --- | --- | --- | --- | --- | --- | --- |
| Pepper | R<sup>2</sup> | Allelic effect | R<sup>2</sup> | Allelic effect | R<sup>2</sup> | Allelic effect | R<sup>2</sup> | Allelic effect |
| Archimedes | 0.03 | -5.89 | 0.01 | -2.53 | 0.05 | -6.77 | 0.03 | -3.72 |
| Aristotle | 0.1 | -17.78 | 0.11 | -11.85 | 0.21 | -22.85 | 0.12 | -11.58 |
| EarlyJalapeno | 0.05 | -6.57 | 0.05 | -4.2 | 0.07 | -7.54 | 0.03 | -3.32 |
| Paladin | 0.04 | -6.38 | 0.04 | -3.91 | 0.05 | -6.12 | 0.04 | -3.59 |
| Perennial | 0.09 | -15.73 | 0.01 | -3.85 | 0.15 | -18.09 | 0.02 | -4.23 |
| RedKnight | 0.14 | -23.77 | 0.28 | -20.67 | 0.19 | -23.76 | 0.28 | -19.98 |
| Revolution | 0.13 | -16.17 | 0.07 | -8.05 | 0.15 | -16.12 | 0.08 | -7.86 |
| Vanguard | 0.09 | -16.56 | 0.06 | -8.63 | 0.12 | -16.48 | 0.09 | -9.85 |
| Across-pepper | 0.13 | -13.64 | 0.11 | -8.07 | 0.19 | -14.74 | 0.13 | -8.04 |
| PhenoPC2 | 0.03 | 0.55 | 0.03 | 0.34 | 0.04 | 0.57 | 0.03 | 0.34 |
| PhenoPC3 | 0.01 | 0.21 | 0.15 | 0.53 | 0.02 | 0.24 | 0.14 | 0.48 |
| PhenoPC4 | 0 | 0.08 | 0.04 | 0.2 | 0.01 | -0.1 | 0.06 | 0.23 |

<sup>a</sup> Models fit using non-transformed phenotypic distributions without any covariates.

**Table S6:**
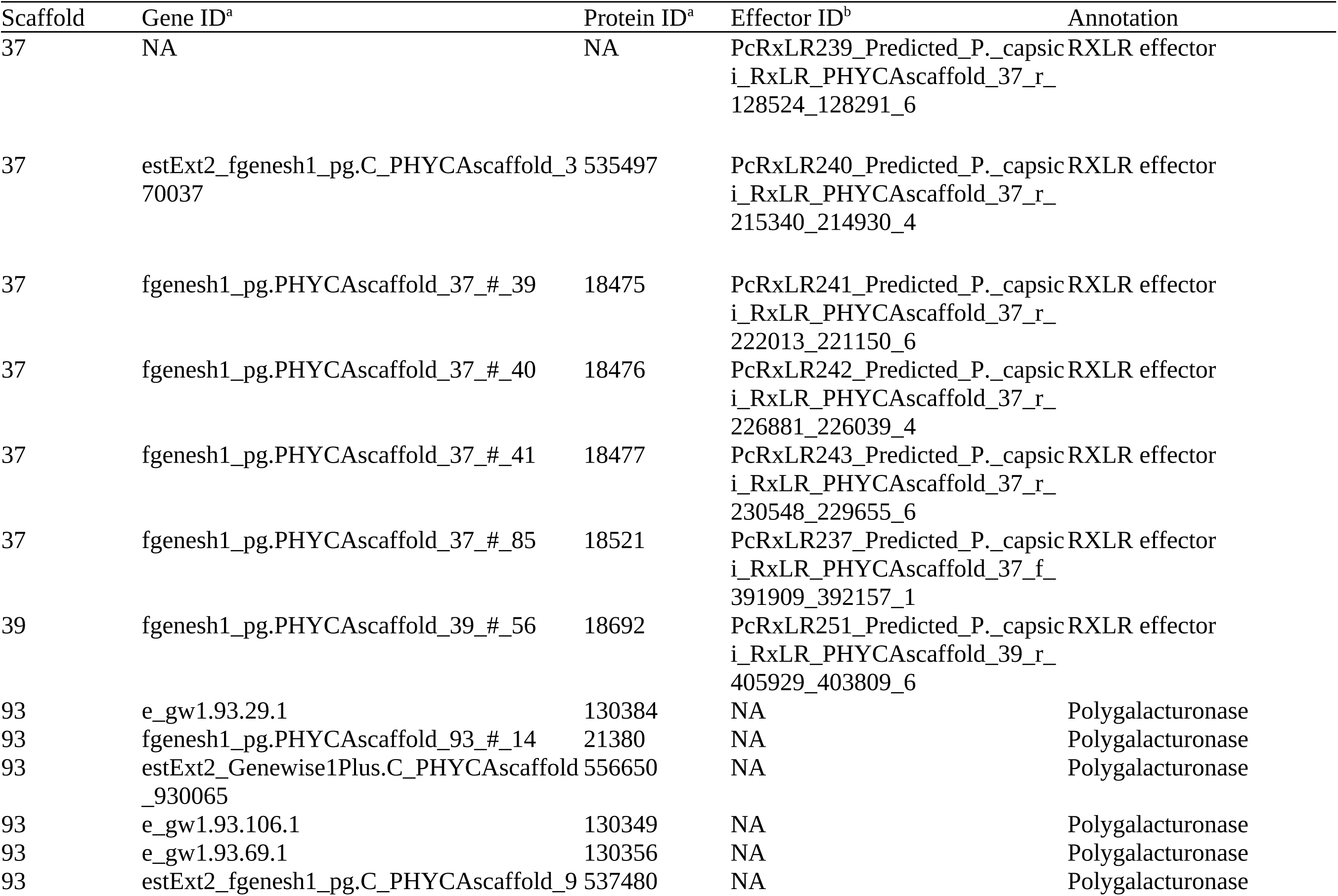
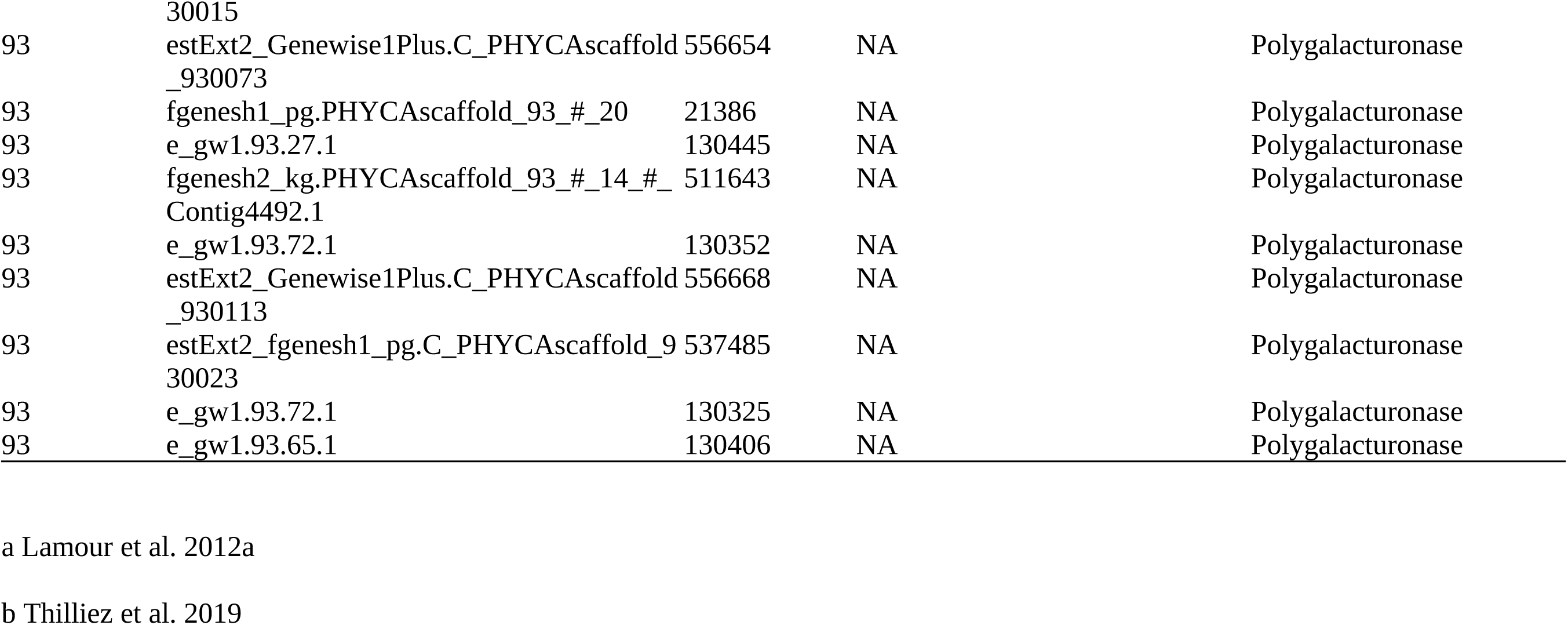
Candidate genes in regions of LD with peak SNPs identified via GWAS.

## SUPPLEMENTAL FIGURE CAPTIONS

**Figure S1:** The percentage of isolates that killed at least one plant in both experimental replicates for the 16 pepper accessions included in the experiment. Numbers above bars refer to the absolute number of isolates deemed pathogenic on that pepper accession. The dashed line at 20% denotes the cutoff used for removing non-informative pepper accessions from the dataset.

**Figure S2:** Scree plots showing A) the variance explained by each principal component from the principal component analysis of the genotype matrix and B) within-groups sum of squares as a function of the number of clusters used in k-means clustering.

**Figure S3:** Loadings of the eight pepper accessions on principal components 1-4 (panes A-D, respectively) from the phenotypic principal component analysis.

**Figure S4:** Manhattan and Q-Q plots showing *P*-values from genome-wide association studies for virulence on each of A) Archimedes, B) Aristotle, C) Early Jalapeño, D) Paladin, and E) Vanguard; and phenotypic PCs 2-4 (F-H).

## Notes

### Competing Interest Statement

The authors have declared no competing interest.

