## Supplementary figures and images for "Quantitative genetic analysis of interactions in the pepper-*Phytophthora capsici* pathosystem"

### Figure S1

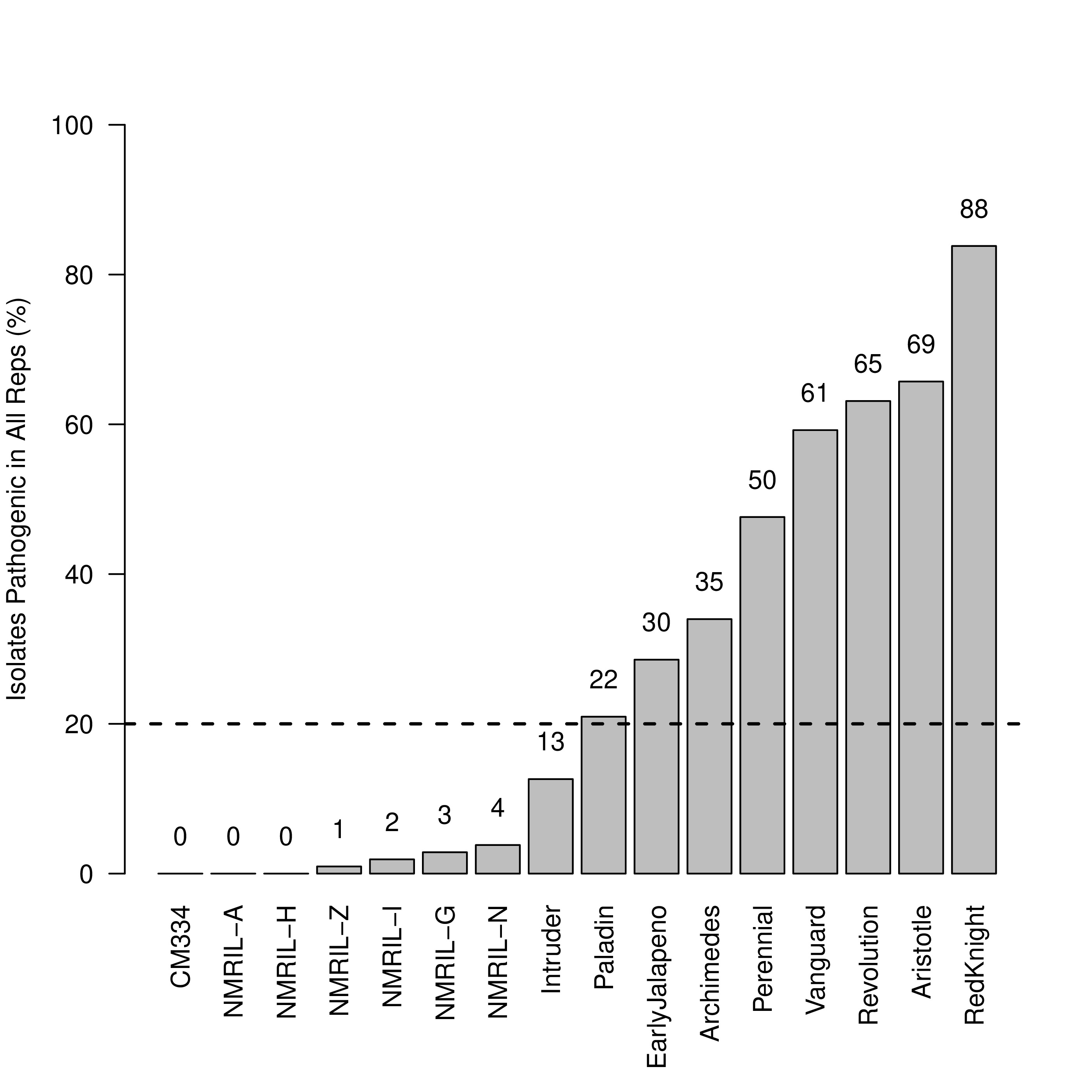

### Figure S2

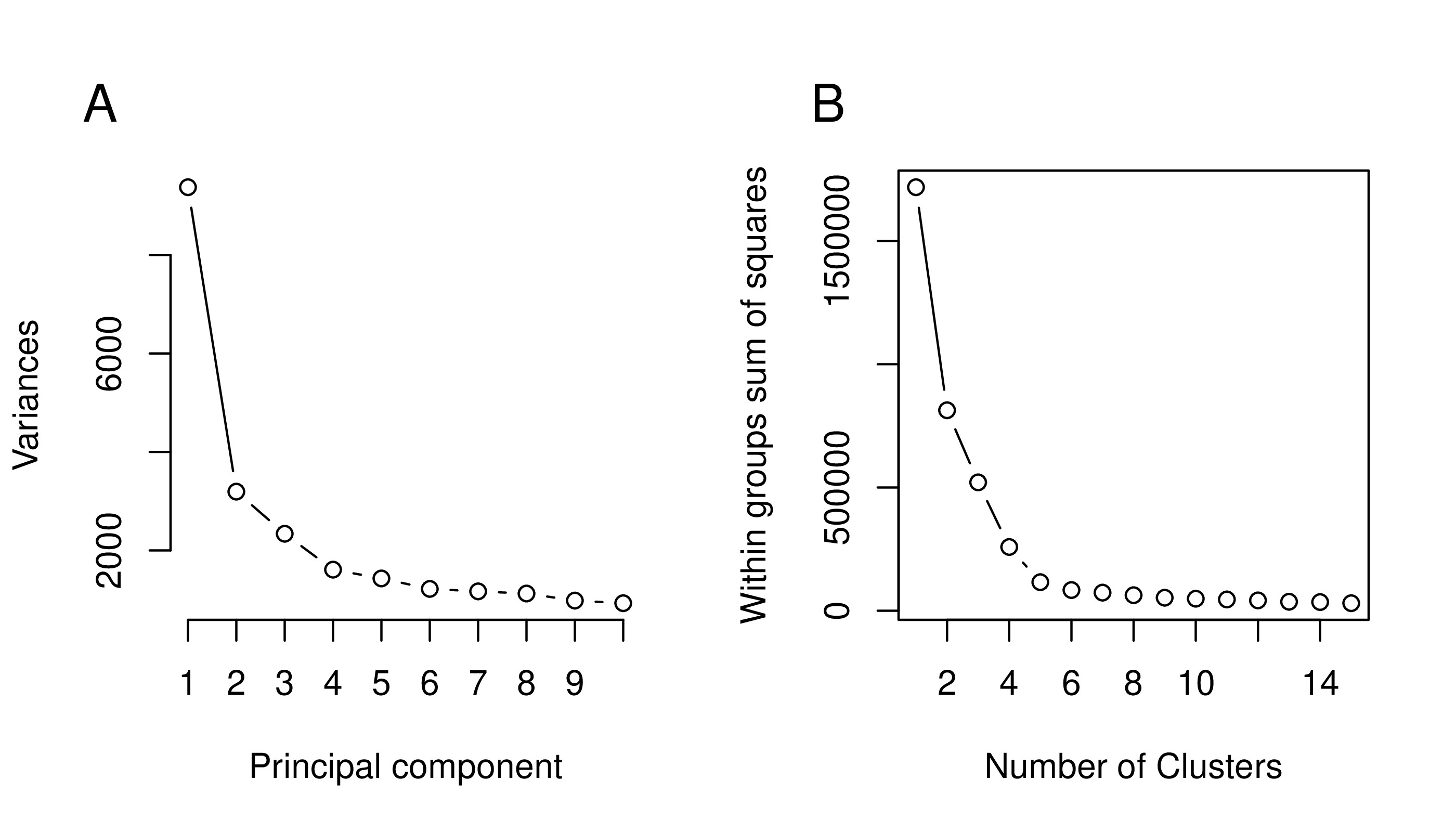

### Figure S3

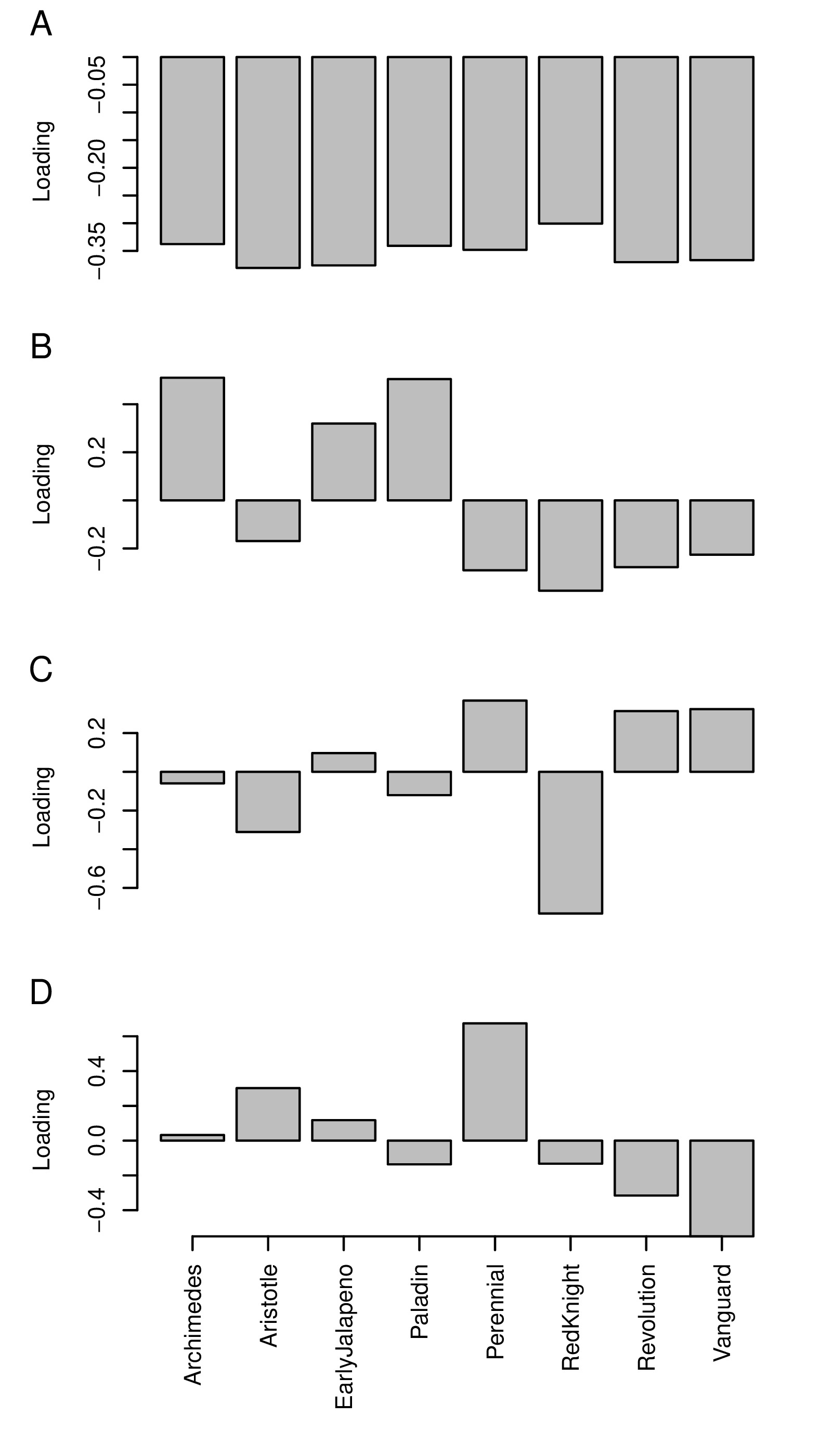

### Figure S4

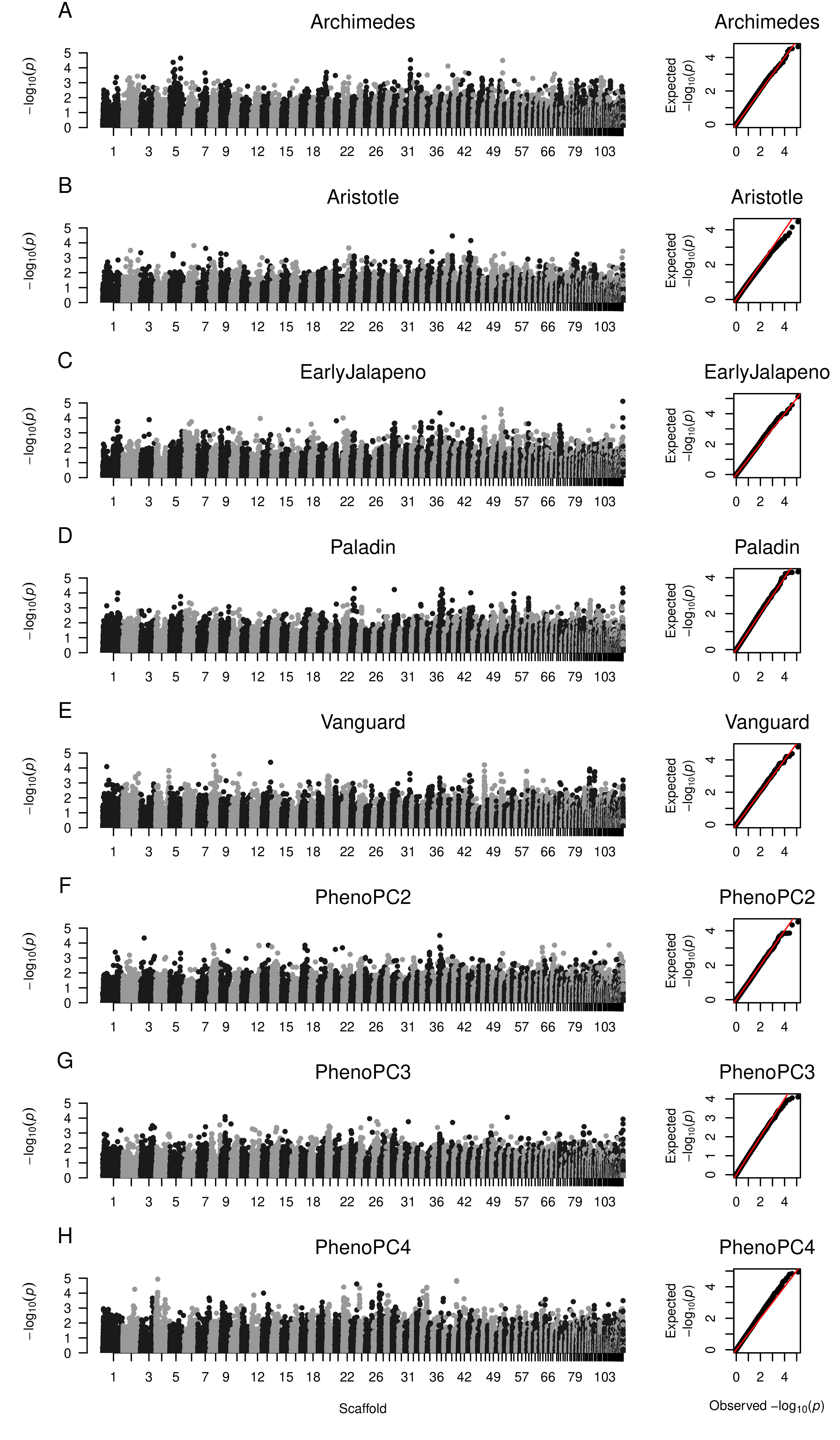
